# SnapATAC: A Comprehensive Analysis Package for Single Cell ATAC-seq

**DOI:** 10.1101/615179

**Authors:** Rongxin Fang, Sebastian Preissl, Yang Li, Xiaomeng Hou, Jacinta Lucero, Xinxin Wang, Amir Motamedi, Andrew K. Shiau, Xinzhu Zhou, Fangming Xie, Eran A. Mukamel, Kai Zhang, Yanxiao Zhang, M. Margarita Behrens, Joseph R. Ecker, Bing Ren

## Abstract

Identification of the *cis*-regulatory elements controlling cell-type specific gene expression patterns is essential for understanding the origin of cellular diversity. Conventional assays to map regulatory elements via open chromatin analysis of primary tissues is hindered by heterogeneity of the samples. Single cell analysis of transposase-accessible chromatin (scATAC-seq) can overcome this limitation. However, the high-level noise of each single cell profile and the large volumes of data could pose unique computational challenges. Here, we introduce SnapATAC, a software package for analyzing scATAC-seq datasets. SnapATAC can efficiently dissect cellular heterogeneity in an unbiased manner and map the trajectories of cellular states. Using the Nyström method, a sampling technique that generates the low rank embedding for large-scale dataset, SnapATAC can process data from up to a million cells. Furthermore, SnapATAC incorporates existing tools into a comprehensive package for analyzing single cell ATAC-seq dataset. As demonstration of its utility, SnapATAC was applied to 55,592 single-nucleus ATAC-seq profiles from the mouse secondary motor cortex. The analysis revealed ∼370,000 candidate regulatory elements in 31 distinct cell populations in this brain region and inferred candidate transcriptional regulators in each of the cell types.

## Introduction

A multicellular organism comprises diverse cell types, each highly specialized to carry out unique functions. Each cell lineage is established during development as a result of tightly regulated spatiotemporal gene expression programs^1^, which are driven in part by sequence-specific transcription factors that interact with *cis*-regulatory sequences in a cell-type specific manner^2^. Thus, identifying the *cis*-elements and their cellular specificity is an essential step towards understanding the developmental programs encoded in the linear genome sequence.

Since the *cis*-regulatory elements are often marked by hypersensitivity to nucleases or transposases when they are active or poised to act, approaches to detect chromatin accessibility, such as ATAC-seq (Assay for Transposase-Accessible Chromatin using sequencing)^3^ and DNase-seq (DNase I hypersensitive sites sequencing)^4^ have been widely used to map candidate *cis*-regulatory sequences. However, conventional assays that use bulk tissue samples as input cannot resolve cell-type specific usage of *cis* elements and lacks the resolution to study their temporal dynamics. To overcome these limitations, a number of methods have been developed for measuring chromatin accessibility in single cells. One approach involves combinatorial indexing to simultaneously analyze tens of thousands of cells^5^. This strategy has been successfully applied to embryonic tissues in *D. melanogaster*^6^, developing mouse forebrains^7^ and adult mouse tissues^8^. A related method, scTHS-seq (single-cell transposome hypersensitive site sequencing), has also been used to study chromatin landscapes at single cell resolution in the adult human brains^9^. A third approach relies on isolation of cell using microfluidic devices (Fluidigm, C1)^10^ or within individually indexable wells of a nano-well array (Takara Bio, ICELL8)^11^. More recently, single cell ATAC-seq analysis has been demonstrated on droplet-based platforms^12,13^, enabling profiling of chromatin accessibility from hundreds of thousands cells in a single experiment^13^. Hereafter, these methods are referred to collectively as single cell ATAC-seq (scATAC-seq).

The growing volume of scATAC-seq datasets coupled with the sparsity of signals in each individual profile due to low detection efficiency (5-15% of peaks detected per cell)^7^ present a unique computational challenge. To address this challenge, a number of unsupervised algorithms have been developed. One approach, chromVAR^14^, groups similar cells together by dissecting the variability of transcription factor (TF) motif occurrence in the open chromatin regions in each cell. Another approach employs the natural language processing techniques such as Latent Semantic Analysis (LSA)^8^ and Latent Dirichlet Allocation (LDA)^15^ to group cells together based on the similarity of chromatin accessibility. A third approach analyzes the variability of chromatin accessibility in cells based on the k-mer composition of the sequencing reads from each cell^13,16^. A fourth approach, Cicero^17^, infers cell-to-cell similarities based on the gene activity scores predicted from their putative regulatory elements in each cell.

Because the current methods often require performing linear dimensionality reduction such as singular value decomposition (SVD) on a cell matrix of hundreds of thousands of dimensions, scaling the analysis to millions of cells remains very challenging or nearly impossible. In addition, the unsupervised identification of cell types or states in complex tissues using scATAC-seq dataset does not have the same degree of sensitivity as that from scRNA-seq^18^. One possibility is that the current methods rely on the use of pre-defined accessibility peaks based on the aggregate signals. There are several limitations to this choice. First, the cell type identification could be biased toward the most abundant cell types in the tissues, and consequently lack the ability to reveal regulatory elements in the rare cell populations that could be underrepresented in the aggregate dataset. Second, a sufficient number of single cell profiles would be required to create robust aggregate signal for creating the peak reference.

To overcome these limitations, we introduce a software package, Single Nucleus Analysis Pipeline for ATAC-seq – SnapATAC (https://github.com/r3fang/SnapATAC) - that does not require population-level peak annotation prior to clustering. Instead, it resolves cellular heterogeneity by directly comparing the similarity in genome-wide accessibility profiles between cells. We also adopt a new sampling technique, ensemble Nyström method^19,20^, that significantly improves the computational efficiency and enables the analysis of scATAC-seq from a million cells on typical hardware. SnapATAC also incorporates many existing tools, including integration of scATAC-seq and scRNA-seq dataset^18^, prediction of enhancer-promoter interaction, discovery of key transcription factors^21^, identification of differentially accessible elements^22^, construction of trajectories during cellular differentiation, correction of batch effect^23^ and classification of new dataset based on existing cell atlas^18^, into one single package to maximize its utility and functionalities. Thus, SnapATAC represents a comprehensive solution for scATAC-seq analysis.

Through extensive benchmarking using both simulated and empirical datasets from diverse tissues and species, we show that SnapATAC outperforms current methods in accuracy, sensitivity, scalability and reproducibility for cell type identification from complex tissues. Furthermore, we demonstrate the utility of SnapATAC by building a high-resolution single cell atlas of the mouse secondary motor cortex. This atlas comprises of ∼370,000 candidate *cis*-regulatory elements across 31 distinct cell types, including rare neuronal cell types that account for less than 0.1% of the total population analyzed. Through motif enrichment analysis, we further infer potential key transcriptional regulators that control cell type specific gene expression programs in the mouse brain.

## Results

### Overview of SnapATAC workflow

A schematic overview of SnapATAC workflow is displayed in **Fig. 1**. SnapATAC first performs pre-processing of sequencing reads including demultiplexing, reads alignments and filtering, duplicate removal and barcode selection using SnapTools (https://github.com/r3fang/SnapTools) (**Supplementary Methods**). The output of this pre-processing step is a “snap” (Single-Nucleus Accessibility Profiles) file (**Supplementary Note 1**) specially formatted for storing single cell ATAC-seq datasets (**Supplementary Fig. 1a**). Users could select high quality single cell ATAC-seq profiles for subsequent analysis based on numbers of unique fragments detected from the cell and percentage of promoter-overlapping fragments^24^.

**Figure 1.**
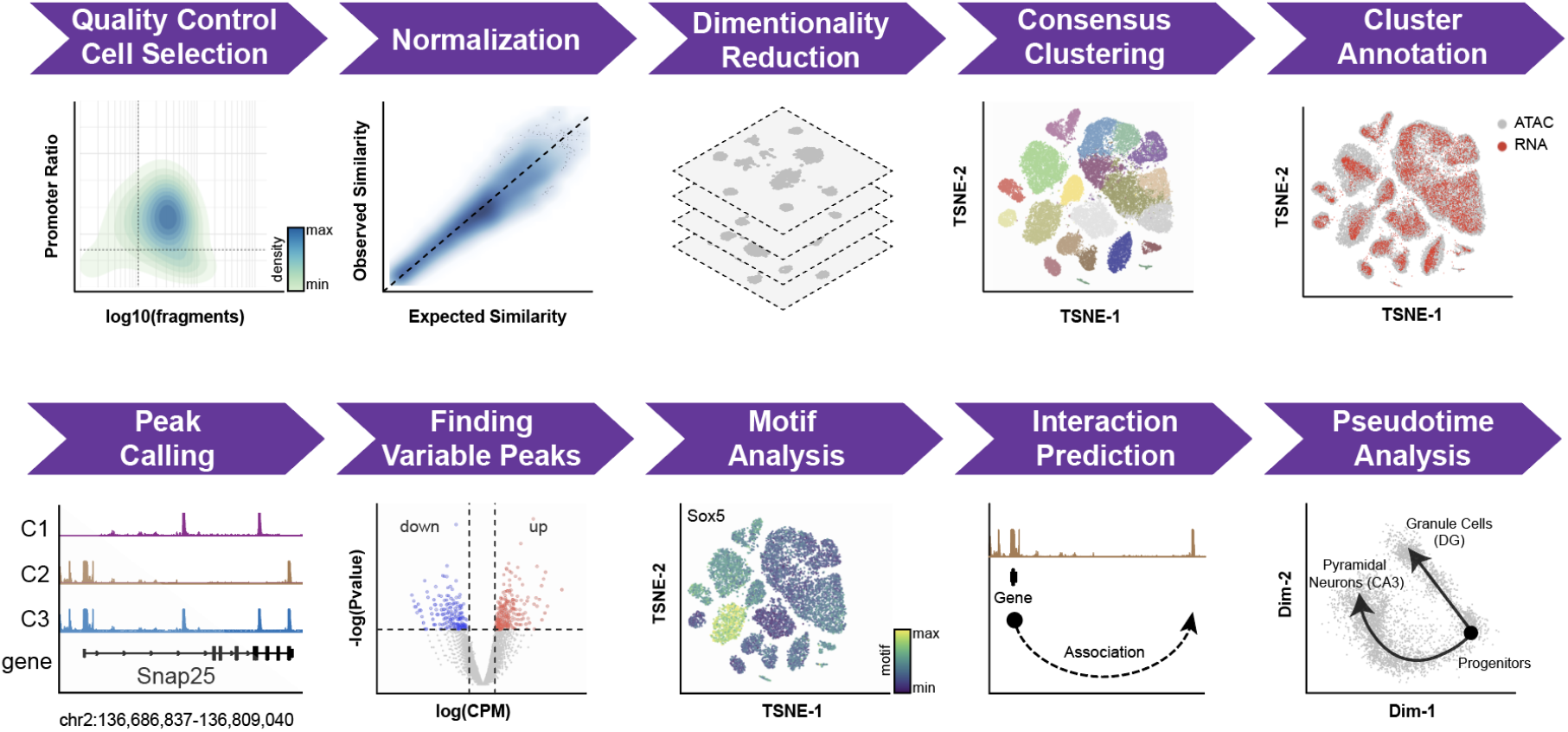
Schematic overview of SnapATAC analysis workflow. See main text for description of each step.

Next, SnapATAC resolves the heterogeneity of cell population by assessing the similarity of chromatin accessibility between cells. To achieve this goal, each single cell chromatin accessibility profile is represented as a binary vector, the length of which corresponds to the number of uniform-sized bins that segment the genome. Through systematic benchmarking, a bin size of 5kb is chosen in this study (**Supplementary Methods** and **Supplementary Fig. 2b**). A bin with value “1” indicates that one or more reads fall within that bin, and the value “0” indicates otherwise. The set of binary vectors from all the cells are converted into a Jaccard similarity matrix, with the value of each element calculated from the fraction of overlapping bins between every pair of cells. Because the value of Jaccard Index could be influenced by sequencing depth of a cell (**Supplementary Methods**), a regression-based normalization method is developed to remove this confounding factor (**Supplementary Methods** and **Supplementary Fig. 3–4**). Using the normalized similarity matrix, eigenvector decomposition is performed for dimensionality reduction. Finally, in the reduced dimension, SnapATAC uses Harmony^23^ to remove potential batch effect between samples introduced by technical variability (**Supplementary Methods**).

The computational cost of the algorithm scales quadratically with the number of cells. To improve the scalability of SnapATAC, a sampling technique - the Nyström method^19^ – is used to efficiently generate the low-rank embedding for large-scale datasets (**Supplementary Methods**). Nyström method contains two major steps: 1) it computes the low dimension embedding for a subset of selected cells (also known as landmarks); 2) it projects the remaining cells to the embedding structure learned from the landmarks. This achieves significant speedup considering that the number of landmarks could be substantially smaller than the total number of cells. Through benchmarking, we further demonstrate that this approach will not sacrifice the performance once the landmarks are chosen appropriately (**Supplementary Methods** and **Supplementary Fig. 5a-c**) as reported before^20^.

Nyström method is stochastic and could yield different clustering results in each sampling. To overcome this limitation, a consensus approach is used that combines a mixture of low-dimensional manifolds learned from different sets of sampling (**Supplementary Methods**). Through benchmarking, we demonstrate that the ensemble approach can significantly improve the reproducibility of clustering outcome compared to the standard Nyström method (**Supplementary Fig. 5d**). In addition, this consensus algorithm naturally fits within the distributed computing environments where their computational costs are roughly the same as that of the standard single sampling method.

As a standalone software package, SnapATAC also provides a number of commonly used functions for scATAC-seq analysis by incorporating many existing useful tools, as described below:

First, to facilitate the annotation of resulting cell clusters, SnapATAC provides three different approaches: i) SnapATAC annotates the clusters based on the accessibility score at the canonical marker genes (**Supplementary Methods**); ii) it infers cell type labels by integrating with corresponding single cell RNA-seq datasets^18^ (**Supplementary Methods** and **Fig. 2a**); iii) it allows supervised annotation of new single cell ATAC-seq dataset based on an existing cell atlas (**Supplementary Methods**).

**Figure 2.**
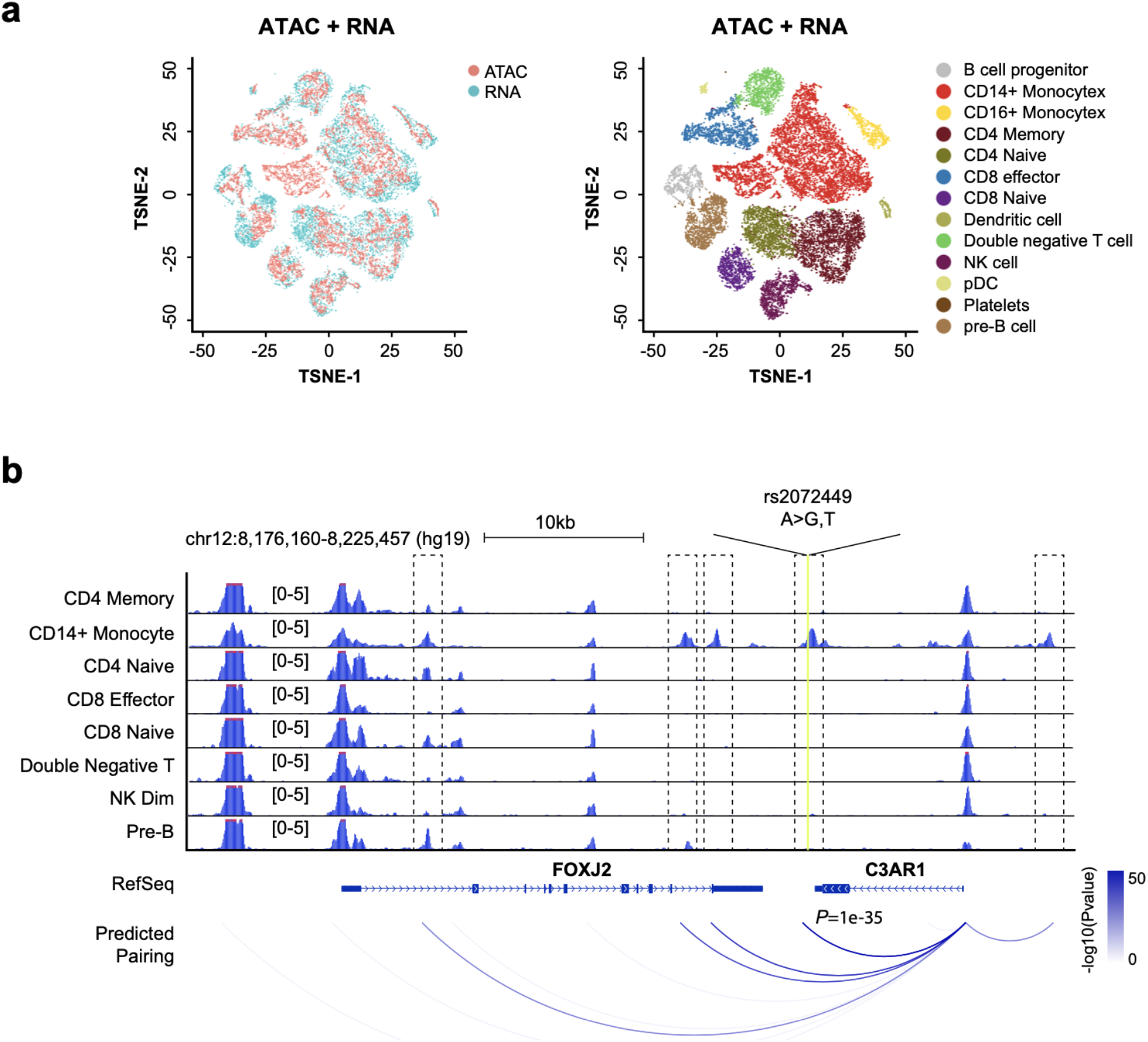
SnapATAC integrates single cell ATAC-seq and RNA-seq data to link enhancers to putative target genes. (**a**) Joint t-SNE visualization of scATAC-seq and scRNA-seq datasets from peripheral blood mononuclear cells (PBMC). Cells are colored by modality (left) and predicted cell types (right). (**b**) Cell-type specific chromatin landscapes are shown together with the association score between gene expression of C3AR1 and accessibility at its putative enhancers. Dash lines highlight the significant enhancer-promoter pairs. Yellow line represents the SNP (rs2072449) that is associated with C3AR1 expression^25^.

Second, SnapATAC allows identification of the candidate regulatory elements in each cluster by applying various peak-calling algorithms^26^ to the aggregate chromatin profiles. Differential analysis is then performed to identify cell-type specific regulatory elements^22^. Candidate master transcription factors in each cell cluster are discovered through motif enrichment analysis of the differentially accessible regions in each cluster^27^. SnapATAC further conducts Genomic Regions Enrichment of Annotation Tool (GREAT)^28^ analysis to identify the biological pathways active in each cell type.

Third, SnapATAC incorporates a new approach to link candidate regulatory elements to their putative target genes. In contrast to previous method^17^ that relies on analysis of co-accessibility of putative enhancers and promoters^29^, SnapATAC infers the linkage based on the association between gene expression and chromatin accessibility in single cells where scRNA-seq data is available (**Supplementary Methods**). First, SnapATAC integrates scATAC-seq and scRNA-seq using Canonical Correlation Analysis (CCA) as described in the previous study^30^. Second, for each scATAC-seq profile, a corresponding gene expression profile is imputed based on the weighted average of its *k*-nearest neighboring cells (i.e. k=15) in the scRNA-seq dataset. A “pseudo” cell is created that contains the information of both chromatin accessibility and gene expression. Finally, logistic regression is performed to quantify the association between the gene expression and binarized accessibility state at putative enhancers (**Supplementary Methods**). This new approach is used to integrate ∼15K peripheral blood mononuclear cells (PBMC) chromatin profiles and ∼10K PBMC transcriptomic profiles (**Fig. 2a**) and represent them in a joint t-SNE embedding space (**Fig. 2a**). Over 98% of the single cell ATAC-seq cells can be confidently assigned to a cell type defined in the scRNA-seq dataset (**Supplementary Fig. 6a**). Enhancer-gene pairs are predicted for 3,000 genes differentially expressed between cell types in PBMC as determined by scRNA-seq using Seurat^18^. The validity of the prediction is supported by two lines of evidence. First, the association score exhibits a distance decay from the TSS, consistent with the distance decay of interaction frequency observed in chromatin conformation study^31^ (**Supplementary Fig. 6b**). Second, the predictions match well with the expression quantitative trait loci (*cis*-eQTLs) derived from interferon-γ and lipopolysaccharide stimulation of monocytes^25^ with reasonable prediction power (AUROC=0.66, AUPRC=0.68; **Supplementary Fig. 6c-d** and **Supplementary Methods**). It is important to note that while statistical association between scATAC-seq and scRNA-seq provides another approach to symmetrically link enhancers to their putative target genes, the predictions require further experimental validation.

Fourth, SnapATAC has incorporated a function to construct cellular trajectories from single cell ATAC-seq. As a demonstration of this feature, SnapATAC is used to analyze a dataset that contains 4,259 cells from the hippocampus in the fetal mouse brain (E18) (**Supplementary Table S1**). Immature granule cells originating in the dentate gyrus give rise to both mature granule cells (DG) and pyramidal neurons (CA3)^33^. Analysis of 4,259 cells reveals a clear branching structure in the first two dimensions (**Fig. 3a**), the pattern of which is remarkably similar to the result previously obtained from single cell transcriptomic analysis^34^. For instance, the DG-specific transcription factor *Prox1* is exclusively accessible in one branch whereas *Neurod6* that is known to be specific to CA3 are accessible in the other branch. Markers of progenitors such as *Hes5* and *Mki67*, however, are differentially accessible before the branching point (**Fig. 3b**). Further using lineage inference tool such as Slingshot^32^, SnapATAC defines the trajectories of cell states for pseudo-time analysis (**Fig. 3a**). These results demonstrate that SnapATAC can also reveal lineage trajectories with high accuracy.

**Figure 3.**
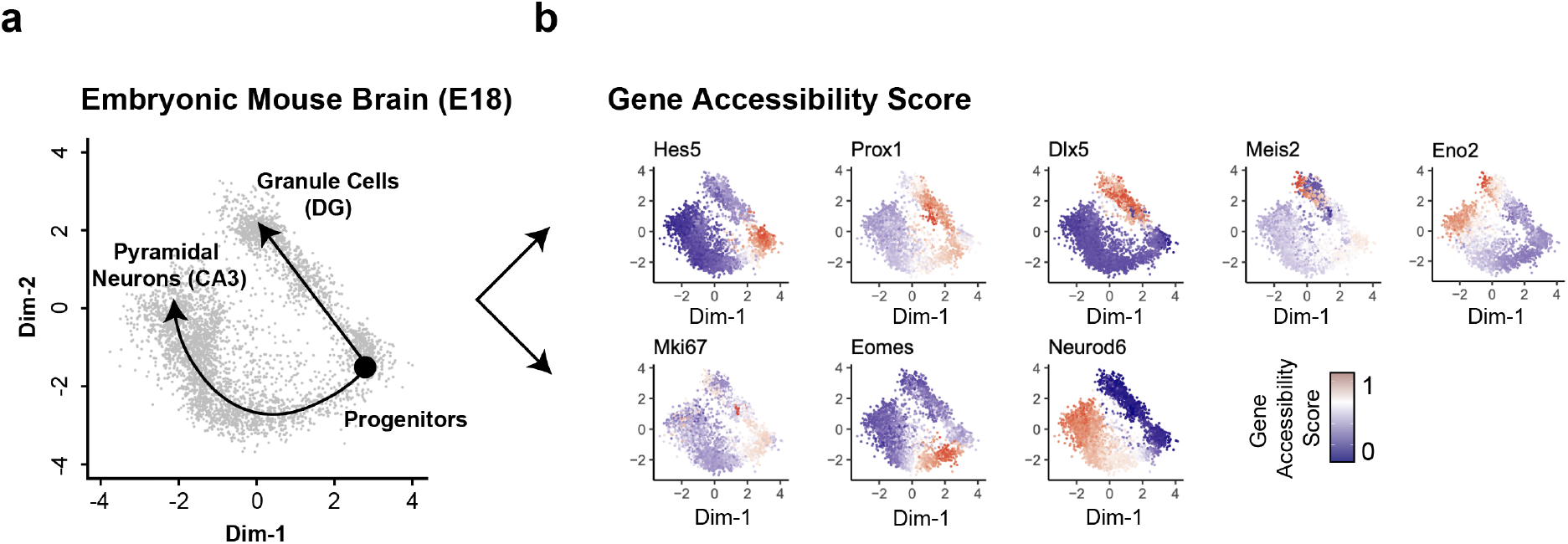
SnapATAC constructs cellular trajectories for the developing mouse brain. (**a**) Two-dimensional visualization of a dataset that contains 4,259 single cell chromatin profiles from the hippocampus and ventricular zone in embryonic mouse brain (E18) reveals two-branch differentiation trajectories from progenitor cells to Granule Cells (DG) and Pyramidal Neurons (CA3) (left). Data source is listed in **Supplementary Table S1**. The cellular trajectory is determined by Slingshot^32^. (**b**) Gene accessibility score of canonical marker genes is projected onto the 2D embedding.

### Performance evaluation

To compare the accuracy of cell clustering between SnapATAC and published scATAC-seq analysis methods, a simulated dataset of scATAC-seq profiles are generated with varying coverages, from 10,000 (high coverage) to 1,000 reads per cell (low coverage) by down sampling from 10 previously published bulk ATAC-seq datasets^27^ (**Supplementary Table S2** and **Supplementary Methods**). Based on a recent summary of cell ATAC-seq methods^35^, LSA^8^ and cisTopic^15^ outperforms the other methods in separating cell populations of different coverages and noise levels in both synthetic and real datasets. Therefore, we choose to compare SnapATAC with these two methods.

The performance of each method in identifying the original cell types is measured by both Adjusted Rank Index (ARI) and Normalized Mutual Index (NMI). The comparison shows that SnapATAC is the most robust and accurate method across all ranges of data sparsity (Wilcoxon signed-rank test, *P* < 0.01; **Fig. 4a**; **Supplementary Fig. 7** and **Supplementary Table S3**). Next, a set of 1,423 human cells corresponding to 10 distinct cell types generated using C1 Fluidigm platform, where the ground truth is known^14^, is analyzed by SnapATAC and other methods. Again, SnapATAC correctly identifies the cell types with high accuracy (**Supplementary Fig. 8**).

**Figure 4.**
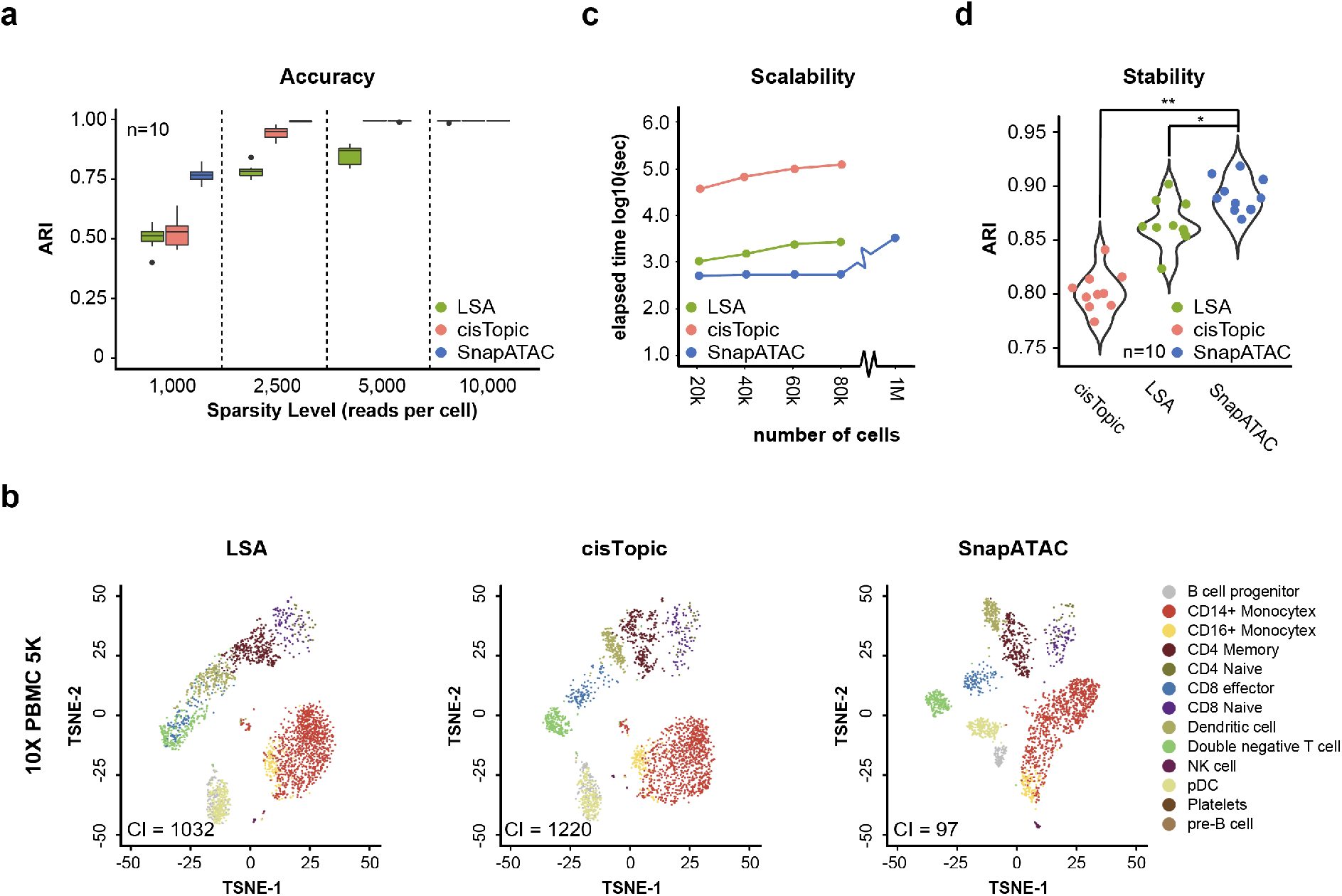
SnapATAC outperforms current methods in accuracy, sensitivity, scalability and stability of identifying cell types in complex tissues. (**a**) A set of simulated datasets are generated with varying coverage ranging from 1,000 to 10,000 reads per cell cells (**Supplementary Methods**). For each coverage, n=10 random replicates are simulated, and clustering accuracy measurement is based on Adjusted Rank Index (ARI). (**b**) T-SNE representation of PBMC single cell ATAC-seq profiles analyzed by LSA (left), cisTopic (middle) and SnapATAC (right). The cell type identification was predicted by 10X PBMC single cell RNA-seq using recent integration method^30^. CI = connectivity index (see **Supplementary Methods**). (**c**) A mouse dataset^8^ is sampled to different number of cells ranging from 20k to 1M. For each sampling, we compared the CPU running time of different methods for dimensionality reduction (**Supplementary Methods**). SnapATAC is the only method that is able to process a dataset of one million (1M) cells. (**d**) A set of perturbations (n=5) are introduced to the mouse atlas dataset by down sampling to 90% of the original sequencing depth. Clustering outcomes are compared between different down sampled datasets (n=10) to estimate the reproducibility. One-tailed t-test was performed to estimate the significance level between SnapATAC and each of the other two methods (* < 0.05 and ** < 0.01).

To compare the sensitivity of SnapATAC on detecting cell types to that of previously published methods, we analyzed two scATAC-seq datasets representing different types of bio-samples. First, to quantify the clustering sensitivity, we applied an existing integration method to predict the cell type of 4,792 PBMC cells using corresponding 10X single cell RNA-seq by following the tutorial (https://satijalab.org/seurat/v3.1/atacseq_integration_vignette.html). To obtain the most confident prediction, we only kept single cell ATAC-seq profiles whose cell type prediction score is greater than 0.9. Using the remaining cells, we calculated the connectivity index (CI; **Supplementary Methods**) in the low-dimension manifold for each of the methods (LSA, cisTopic and SnapATAC). Connectivity index estimates the degree of separation between clusters in an unbiased manner and a lower connectivity index represents a higher degree of separation between clusters. SnapATAC exhibits substantially higher sensitivity in distinguishing different cell types compared to the other two methods (**Fig. 4b**). The second is a newly produced dataset that contains 9,529 single nucleus open chromatin profiles generated from the mouse secondary motor cortex. Based on the gene accessibility score at canonical marker genes (**Supplementary Fig. 9**), SnapATAC uncovers 22 distinct cell populations (**Supplementary Fig. 10**) whereas alternative methods fail to distinguish the rare neuronal subtypes including Sst (Gad2+ and Sst+), Vip (Gad2+ and Vip+), L6b (Sulf1- and Tl4e+) and L6.CT (Sulf1+ and Foxp2+). These results suggest that SnapATAC outperforms existing methods in sensitivity of separating different cell types in both synthetic and real datasets.

To compare the scalability of SnapATAC to that of existing methods, a previous scATAC-seq dataset that contains over 80k cells from 13 different mouse tissues^8^ is used (**Supplementary Table S1**). This dataset is down sampled to different number of cells, ranging from 20,000 to 80,000 cells. For each sampling, SnapATAC and other methods are performed, and the CPU running time of dimensionality reduction is monitored (**Supplementary Methods**). The running time of SnapATAC scales linearly and increases at a significantly lower slope than alternative methods (**Fig. 4c**). Using the same computing resource, when applied to 100k cells, SnapATAC is much faster than existing methods (**Fig. 4c**). For instance, when applied to 100k cells, SnapATAC is nearly 10 times faster than LSA and more than 100 times faster than cisTopic. More importantly, because SnapATAC avoids the loading of the full cell matrix in the memory and can naturally fit within the distributed computing environments (**Supplementary Methods**), the running time and memory usage for SnapATAC plateau after 20,000 cells, making it possible for analyzing datasets of even greater volumes. To test this, we simulate one million cells of the same coverage with the above dataset (**Supplementary Methods**) and process it with SnapATAC, LSA and cisTopic. Using the same computing resource, SnapATAC is the only method that is able to process this dataset (**Fig. 4c** and **Supplementary Methods**). These results demonstrate that SnapATAC provides a highly scalable approach for analyzing large-scale scATAC-seq dataset.

To evaluate the clustering reproducibility, the above mouse scATAC-seq dataset is down-sampled to 90% of the original sequencing depth in five different iterations. Each down sampled dataset is clustered using SnapATAC and other methods. Clustering results are compared between sampled datasets to estimate the stability. SnapATAC has a substantially higher reproducibility of clustering results between different down-sampled datasets than other methods (**Fig. 4d**).

The improved performance of SnapATAC likely results from the fact that it considers all reads from each cell, not just the fraction of reads within the peaks defined in the population. To test this hypothesis, clustering is performed after removing the reads overlapping the predefined peak regions. Although the outcome is worse than the full dataset as expected, it still recapitulates the major cell types obtained from the full dataset (**Supplementary Fig. 11**). This holds true for all three datasets tested (**Supplementary Fig. 11a-c**). One possibility is that the off-peak reads may be enriched for the euchromatin (or compartment A) that strongly correlates with active genes^28^ and varies considerably between cell types^29,30^. Consistent with this hypothesis, the density of the non-peak reads in scATAC-seq library is highly enriched for the euchromatin (compartment A) as defined using genome-wide chromatin conformation capture analysis (i.e. Hi-C) in the same cell type^31^ (**Supplementary Fig. 12**). These observations suggest that the non-peak reads discarded by existing methods can actually contribute to distinguish different cell types.

Including the off-peak reads, however, raises a concern regarding whether SnapATAC is sensitive to technical variations (also known as batch effect). To test this, SnapATAC is applied to four datasets generated using different technologies (**Supplementary Table S1**). Each dataset contains at least two biological replicates produced by the same technology. In all cases, the biological replicates are well mixed in the t-SNE embedding space showing no batch effect (**Supplementary Fig. 13**), suggesting that SnapATAC is robust to the technical variations.

To test whether SnapATAC is robust to technical variation introduced by different technological platforms, it is used to integrate two mouse brain datasets generated using plate and droplet-based scATAC-seq technologies (**Supplementary Table S1**). In the joint t-TSNE embedding space, these two datasets are separated based on the technologies (**Supplementary Fig. 14a**). To remove the platform-to-platform variations, Harmony^23^, a single cell batch effect correction tool, is incorporated into the SnapATAC pipeline (**Supplementary Methods**). After applying Harmony^23^, these two datasets are fully mixed in the joint t-SNE embedding (**Supplementary Fig. 14b**) and clusters are fairly represented by both datasets (**Supplementary Fig. 14c**).

### A high-resolution *cis*-regulatory atlas of the mouse secondary motor cortex

To demonstrate the utility of SnapATAC in resolving cellular heterogeneity of complex tissues and identify candidate *cis*-regulatory elements in diverse cell type, it is applied to a new single nucleus ATAC-seq dataset generated from the secondary mouse motor cortex in the adult mouse brain as part of the BRAIN Initiative Cell Census Consortium^36^ (**Supplementary Fig. 15a**). This dataset includes two biological replicates, each pooled from 15 mice to minimize potential batch effects. The aggregate signals show high reproducibility between biological replicates (Pearson correlation = 0.99; **Supplementary Fig. 15b-d**) and a significant enrichment for transcription start sites (TSS), indicating a high signal-to-noise ratio (**Supplementary Fig. 15e**). After filtering out the low-quality nuclei (**Supplementary Fig. 16a**) and removing putative doublets using Scrublet^37^ (**Supplementary Methods**; **Supplementary Fig. 16b**), a total of 55,592 nuclear profiles with an average of ∼5,000 unique fragments per nucleus remain and are used for further analysis (**Supplementary Table S4**). To our knowledge, this dataset represents one of the largest single cell chromatin accessibility studies for a single mammalian brain region to date.

SnapATAC identifies initially a total of 20 major clusters using the consensus clustering approach (**Supplementary Fig. 17**). The clustering result is highly reproducible between biological replicates (Pearson correlation=0.99; **Supplementary Fig. 18a**) and is resistant to sequencing depth effect (**Supplementary Fig. 18b**). Based on the gene accessibility score at the canonical marker genes (**Supplementary Fig. 19**), these clusters are classified into 10 excitatory neuronal subpopulations (Snap25+, Slc17a7+, Gad2-; 52% of total nuclei), three inhibitory neuronal subpopulations (Snap25+, Gad2+; 10% of total nuclei), one oligodendrocyte subpopulation (Mog+; 8% of total nuclei), one oligodendrocyte precursor subpopulation (Pdgfra+; 4% of total nuclei), one microglia subpopulation (C1qb+; 5% of total nuclei), one astrocyte subpopulation (Apoe+; 12% of total nuclei), and additional populations of endothelial, and smooth muscle cells accounting for 6% of total nuclei (**Fig. 5a**).

**Figure 5.**
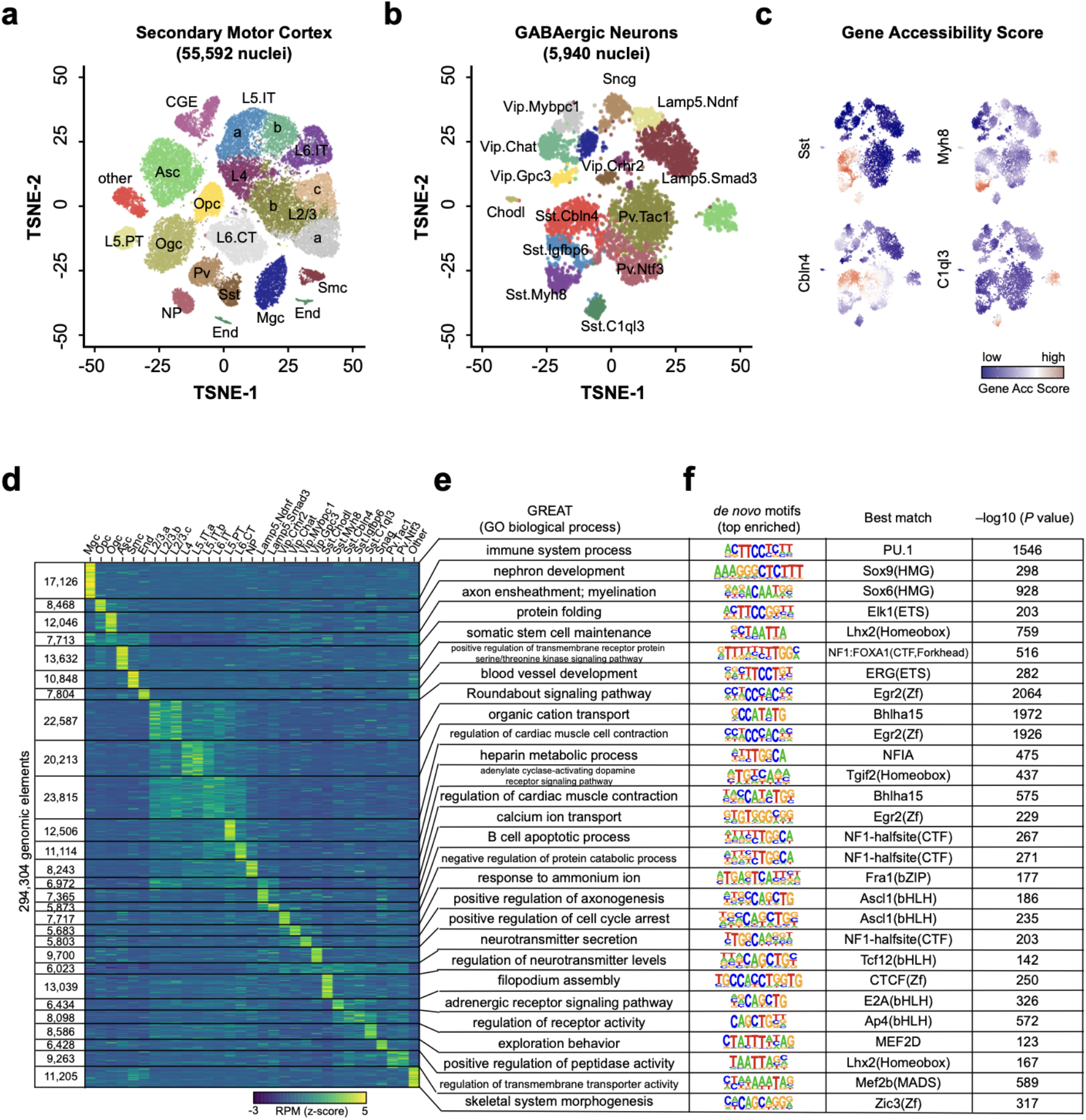
A high-resolution *cis*-regulatory atlas of mouse secondary motor cortex (MOs). (**a**) T-SNE visualization of 20 cell types in MOs identified using SnapATAC. (**b**) Fourteen GABAergic subtypes revealed by iterative clustering of 5,940 GABAergic neurons (Sst, Pv and CGE). (**c**) Gene accessibility score of canonical marker genes for GABAergic subtypes projected onto the t-SNE embedding. Marker genes were identified from previous scRNA-seq analysis^38^. (**d**) *k*-means clustering of 294,304 differentially accessible elements based on chromatin accessibility. (**e**) Gene ontology analysis of each cell type predicted using GREAT analysis^39^. (**f**) Transcription factor motif enriched in each cell group identified using Homer^21^.

In mammalian brain, GABAergic interneurons exhibit spectacular diversity that shapes the spatiotemporal dynamics of neural circuits underlying cognition^40^. To examine whether iterative analysis could help tease out various subtypes of GABAergic neurons, SnapATAC is applied to the 5,940 GABAergic nuclei (CGE, Sst and Vip) identified above, finding 17 distinct sub-populations (**Supplementary Fig. 20a**) that are highly reproducible between biological replicates (Pearson correlation = 0.99; **Supplementary Fig. 20b**). Based on the chromatin accessibility at the marker genes (**Supplementary Fig. 21**), these 17 clusters are classified into five Sst subtypes (Chodl+, Cbln4+, Igfbp6+, Myh8+ and C1ql3+), two Pv subtypes (Tac1+ and Ntf3+), two Lamp5 subtypes (Smad3+ and Ndnf+), four Vip subtypes (Mybpc1+, Chat+, Gpc3+, Crhr2+), Sncg and putative doublets (**Fig. 5b**). These clusters include a rare type Sst-Chodl (0.1%) previously identified in single cell RNA^38^ and single cell ATAC-seq analysis^41^. While the identity and function of these subtypes require further experimental validation, our results demonstrate the exquisite sensitivity of SnapATAC in resolving distinct neuronal subtypes with only subtle differences in the chromatin landscape.

A key utility of single cell chromatin accessibility analysis is to identify regulatory sequences in the genome. By pooling reads from nuclei in each major cluster (**Fig. 5a**), cell-type specific chromatin landscapes can be obtained (**Supplementary Fig. 22** and **Supplementary Methods**). Peaks are determined in each cell type, resulting in a total of 373,583 unique candidate *cis*-regulatory elements. Most notably, 56% (212,730/373,583) of these open chromatin regions cannot be detected from bulk ATAC-seq data of the same brain region (**Supplementary Methods**). The validity of these additional open chromatin regions identified from scATAC-seq data are supported by several lines of evidence. First, these open chromatin regions are only accessible in minor cell populations (**Supplementary Fig. 23a**) that are undetectable in the bulk ATAC-seq signal. Second, these sequences show significantly higher conservation than randomly selected genomic sequences with comparable mappability scores (**Supplementary Fig. 23c**). Third, these open chromatin regions display an enrichment for transcription factor (TF) binding motifs corresponding to the TFs that play important regulatory roles in the corresponding cell types. For example, the binding motif for Mef2c is highly enriched in novel candidate *cis*-elements identified from Pvalb neuronal subtype (P-value = 1e-363; **Supplementary Fig. 23d**), consistent with previous report that Mef2c is upregulated in embryonic precursors of Pv interneurons^42^. Finally, the new open chromatin regions tend to test positive in transgenic reporter assays. Comparison to the VISTA enhancer database^43^ shows that enhancer activities of 256 of the newly identified open chromatin regions have been previously tested using transgenic reporter assays in e11.5 mouse embryos. Sixty five percent (167/256; 65%) of them drive reproducible reporter expression in at least one embryonic tissue, which was substantially higher than background rates (9.7%) estimated from regions in the VISTA database that lack canonical enhancer mark^44^. Four examples are displayed (**Supplementary Fig. 23e**).

SnapATAC identifies 294,304 differentially accessible elements between cell types (**Supplementary Methods**; **Fig. 5d**). GREAT analysis (**Fig. 5e**) and motif inference (**Fig. 5f**) identify the master regulators and transcriptional pathways active in each of the cell types. For instance, the binding motif for ETS-factor PU.1 is highly enriched in microglia-specific candidate CREs, motifs for SOX proteins are enriched in Ogc-specific elements, and bHLH motifs are enriched in excitatory neurons-specific CREs (**Fig. 5f**). Interestingly, motifs for candidate transcriptional regulators, including NUCLEAR FACTOR 1 (NF1), are also enriched in candidate CREs detected in two inhibitory neuron subtypes (Lamp5.Ndnf and Lamp5.Smad3). Motif for CTCF, a multifunctional protein in genome organization and gene regulation^45^, is highly enriched in Sst-Chodl, indicating that CTCF may play a role in neurogenesis. Finally, motifs for different basic-helix-loop-helix (bHLH) family transcription factors, known determinants of neural differentiation^46^, show enrichment for distinct Sst subtypes. For instance, E2A motif is enriched in candidate CREs found in Sst.Myh8 whereas AP4 motif is specifically enriched in peaks found in Sst.Cbln4, suggesting specific role that different bHLH factors might play in different neuronal subtypes.

### SnapATAC enables reference-based annotation of new scATAC-seq datasets

Unsupervised clustering of scATAC-seq datasets frequently requires manual annotation, which is labor-intensive and limited to prior knowledge. To overcome this limitation, SnapATAC provides a function to project new single cell ATAC-seq datasets to an existing cell atlas to allow for supervised annotation of cells. First, the Nystrom method is used to project the query cells to the low-dimension manifold pre-computed from the reference cells (**Supplementary Methods**). In the joint manifold, a neighborhood-based classifier is used to determine the cell type of each query cell based on the label of its *k* nearest neighboring cells in the reference dataset (**Supplementary Methods**). The accuracy of this method is determined by five-fold cross validation using the mouse motor cortex atlas. On average, 98% (±1%) of the cells can be correctly classified, suggesting a high accuracy of the method (**Fig. 6a**).

**Figure 6.**
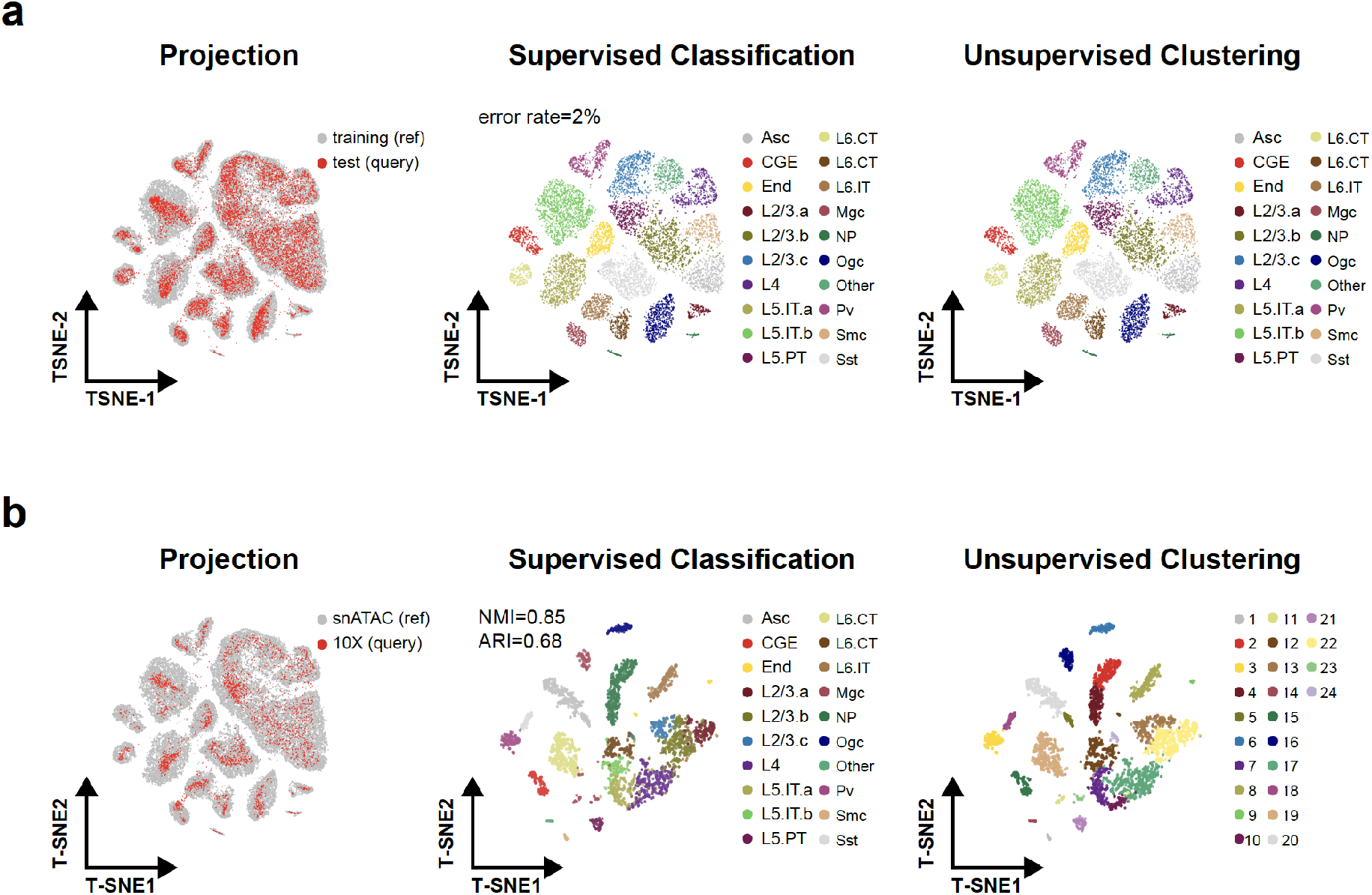
SnapATAC enables supervised annotation of new scATAC-seq dataset using reference cell atlas. (**a**) MOs snATAC-seq dataset is split into 80% and 20% as training and test dataset. A predictive model learned from the training dataset predicts cell types on the test dataset of high accuracy (error rate = 2%) as compared to the original cell type labels (right). (**b**) A predictive model learned from the reference dataset - MOs (snATAC) – accurately predicts the cell types on a query dataset from mouse brain generated using a different technological platform, the 10X scATAC-seq. The t-SNE embedding is inferred from the reference cell atlas (left) or generated by SnapATAC in an unbiased manner from 10X mouse brain dataset (middle and right). Cells are visualized using t-SNE and are colored by the cell types predicted by supervised classification (middle) compared to the cluster labels defined using unsupervised clustering (right).

To demonstrate that SnapATAC could be applied to datasets generated from distinct technical platforms, it is used to annotate 4,098 scATAC-seq profiles from mouse brain cells generated using a droplet-based platform (**Supplementary Table 2**). After removing batch effect introduced by different platforms using Harmony^23^, the query cells are well mixed with the reference cells in the joint embedding space (**Supplementary Fig. 24**). The predicted cluster labels are also consistent with the cell types defined using unbiased clustering analysis (NMI=0.85, ARI=0.68; **Fig. 6b**).

To investigate whether SnapATAC could recognize cell types in the query dataset that are not present in the reference atlas, multiple query data sets are sampled from the above mouse motor cortex dataset and a perturbation is introduced to each sampling by randomly dropping a cell cluster. When this resulting query dataset is analyzed by SnapATAC against the original cell atlas, the majority of the cells that are left out from the original atlas are filtered out due to the low prediction score (**Supplementary Fig. 25**), again suggesting that our method is not only accurate but also robust to the novel cell types in the query dataset.

## Discussion

In summary, SnapATAC is a comprehensive bioinformatic solution for single cell ATAC-seq analysis. The open-source software runs on standard hardware, making it accessible to a broad spectrum of researchers. Through extensive benchmarking, we have demonstrated that SnapATAC outperforms existing tools in sensitivity, accuracy, scalability and robustness of identifying cell types in complex tissues.

SnapATAC differs from previous methods in at least seven aspects. First, SnapATAC incorporates many useful tools and represents the most comprehensive solution for single cell ATAC-seq data analysis to date. In addition to clustering analysis, SnapATAC provides preprocessing, annotation, trajectory analysis, peak calling^26^, differential analysis^22^, batch effect correction^23^ and motif discovery^27^ all in one package. Second, SnapATAC identifies cell types in an unbiased manner without the need for population-level peak annotation, leading to superior sensitivity for identifying rare cell types in complex tissues. Third, SnapATAC utilizes a new algorithm for dimensionality reduction and to identify cell types in heterogeneous tissues and map cellular trajectories. Fourth, with Nyström sampling method^47^, SnapATAC significantly reduces both CPU and memory usage, enabling analysis of large-scale dataset of a million cells or more. Fifth, SnapATAC not only incorporates existing method to integrate scATAC-seq with scRNA-seq dataset^30^ but also provides a new method to predict promoter-enhancer pairing relations based on the statistical association between gene expression and chromatin accessibility in single cells. Sixth, our method achieves high clustering reproducibility using a consensus clustering approach. Finally, SnapATAC also enables supervised annotation of a new scATAC-seq dataset based on an existing reference cell atlas.

It is important to note that a different strategy has been used to overcome the bias introduced by population-based peak annotation^8^. This approach involves iterative clustering, with the first round defining the “crude” clusters in complex tissues followed by identifying peaks in these clusters, which are then used in subsequent round(s) of clustering. However, several limitations still exist. First, the strategy of iterative clustering requires multiple rounds of clustering, aggregation, and peak calling, thus hindering its application to large-scale datasets. Second, the “crude” clusters represent the most dominant cell types in the tissues; therefore, peaks in the rare populations may still be underrepresented. Finally, peak-based methods hinder multi-sample integrative analysis where each sample has its own unique peak reference.

Finally, SnapATAC is applied to a newly generated scATAC-seq dataset including 55,592 high quality single nucleus ATAC-seq profiles from the mouse secondary motor cortex, resulting in a single cell atlas consisting of >370,000 candidate *cis*-regulatory elements across 31 cell types in this mouse brain region. The cellular diversity identified by chromatin accessibility is at an unprecedented resolution and is consistent with mouse neurogenesis and taxonomy revealed by single cell transcriptome data^38,48^. Besides characterizing the constituent cell types, SnapATAC identifies candidate *cis*-regulatory sequences in each of the major cell types and infers the likely transcription factors that regulate cell-type specific gene expression programs. Importantly, a large fraction (56%) of the candidate *cis*-elements identified from the scATAC-seq data are not detected in bulk analysis. While further experiments to thoroughly validate the function of these additional open chromatin regions are needed, the ability for SnapATAC to uncover *cis*-elements from rare cell types of a complex tissue will certainly help expand the catalog of *cis*-regulatory sequences in the genome.

## Data availability

Raw and processed data to support the findings of this study have been deposited to NCBI Gene Expression Omnibus with the accession number GSE126724 with the token of srkxoisclpkppcd.

## Code availability

The scripts and pipeline for the analysis can be found at https://github.com/r3fang/SnapATAC.

## Acknowledgements

We thank D. Gorkin, R. Raviram, and J. Hocker for proofreading and suggestions for the manuscript. We thank S. Kuan for sequencing support. We thank C. Zhang and B. Li for Bioinformatics support. We thank C. O’Connor and C. Fitzpatrick at Salk Institute Flow Cytometry Core for sorting of nuclei. This study was funded by U19MH114831.

## Author Contributions

This study was conceived and designed by R.F. and B.R.; Pipeline developed by R.F.; Data analysis performed by R.F.; Tissue collection and nuclei preparation performed by J.L. and M.B.; Single nucleus ATAC-seq experiment performed by S.P., X.H. and X.W.; Tn5 enzymes synthesized and provided by A.M. and A.S.; Manuscript written by R.F. and B.R. with input from all authors.

## Supplementary Materials

### Outline of the SnapATAC Pipeline

#### Barcode Demultiplexing

Using a custom python script, we first de-multicomplex FASTQ files by integrating the cell barcode into the read name in the following format:

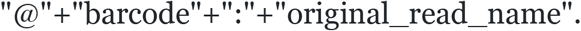

#### Alignment & sorting

Demultiplexed reads are aligned to the corresponding reference genome (i.e. mm10 or hg19) using bwa (0.7.13-r1126) in pair-end mode with default parameter settings. Aligned reads are then sorted based on the read name using samtools (v1.9) to group together reads originating from the same barcodes.

#### Quality Control & Filtering

Pair-end reads are converted into fragments and only those that meet the following criteria are kept: 1) properly paired (according to SMA flag value); 2) uniquely mapped (MAPQ > 30); 3) insert distance within [50-1000bp]. PCR duplicates (fragments sharing exactly the same genomic coordinates) are removed for each cell separately. Given that Tn5 introduces a 9 bp staggered, reads mapping to the positive and negative strand were shifted by +4 / −5bp respectively^49^.

We identify the high-quality cells based on two criteria: 1) total number of unique fragment count [>1,000]; 2) fragments in promoter ratio – the percentage of fragments overlapping with annotated promoter regions [0.2-0.8]. The promoter regions used in this study are downloaded from 10X genomics for hg19 and mm10.

#### Snap File Generation

Using the remaining fragments, we next generate a snap-format (<u>S</u>ingle-<u>N</u>ucleus <u>A</u>ccessibility <u>P</u>rofiles) file using snaptools (https://github.com/r3fang/SnapTools). A snap file is a hierarchically structured hdf5 file that contains the following sections: header (HD), cell-by-bin matrix (BM), cell-by-peak matrix (PM), cell-by-gene matrix (GM), barcode (BD) and fragment (FM). HD session contains snap-file version, date,alignment and reference genome information. BD session contains all unique barcodes and corresponding meta data. BM session contains cell-by-bin matrices of different resolutions. PM session contains cell-by-peak count matrix. GM session contains cell-by-gene count matrix. FM session contains all usable fragments for each cell. Fragments are indexed based on barcodes that enables fast retrieval of reads based on the barcodes. Detailed information about snap file can be found in **Supplementary Note 1**.

##### Box1. Generating Snap file using snaptools

snaptools snap-pre

--input-file=demo.srt.bed.gz
--output-snap=demo.snap
--genome-name=mm10
--genome-size=mm10.gs
--min-mapq=30
--min-flen=50
--max-flen=1000
--keep-single=False
--keep-secondary=False
--keep-discordant=False
--min-cov=0
--max-num=20000
--keep-chrm=True
--overwrite=True

One major utility of the snap file and snaptools is to retrieve reads belonging to a certain group of barcodes. This can be done using snaptools with following command where “barcodes.sel.txt” is a text file that contains the selected barcodes.

##### Box2. Extracting reads using SnapTools

snaptools dump-fragment

--snap-file=demo.snap
--barcode-file=barcodes.sel.txt
--output-file=demo.sel.bed.gz

#### Creating Cell-by-Bin Count Matrix

Using the resulting snap file, we next create cell-by-bin count matrix. The genome is segmented into uniform-sized bins and single cell ATAC-seq profiles are represented as cell-by-bin matrix with each element indicating number of sequencing fragments overlapping with a given bin in a certain cell. In the below example, a cell-by-bin matrix of 5kb resolution is added to demo.snap file.

##### Box 3. Generating cell-by-bin matrix using SnapTools

snaptools snap-add-bmat --snap-file=demo.snap --bin-size-list 5000

#### Optimizing the Bin Size

To evaluate the effect of bin size to clustering performance, we apply SnapATAC to three datasets namely 5K PBMC (10X), Mouse Brain (10X) and MOs-M1 (snATAC). These datasets are generated by both plate and droplet platforms using either cell or nuclei with considerably different depth, allowing us to systematically evaluate the effect of bin size.

For each dataset, we first define the “landmark” cell types in a supervised manner. First, we perform cisTopic^15^ for dimensionality reduction and identify cell clusters using graph-based algorithm Louvain^50^ with k=15. Second, we manually define the major cell types in each dataset by examining the gene accessibility score at the canonical marker genes (**see Supplementary Fig. 9** as an example for MOs-M1). Third, clusters sharing the same marker genes are manually merged and those failing to show unique signatures are discarded. In total, we define nine cell types in PBMC 5K (10X), 14 types in Mouse Brain 5K (10X) and 14 types in MOs M1 (snATAC). Among these cell types, 14 cell populations that account for less than 2% of the total population are considered as rare cell populations (**Supplementary Fig. 2a**).

We next evaluate the performance of each bin size selection using three metrics: 1) cluster connectivity index (CI) which estimate the degree of connectedness of the landmark cell types; a lower CI represents a better separation. The connectivity index is computed in the following manner. For each cell *i,* the K (K=15) nearest neighbors are found and sorted from the closest to furthest. The algorithm checks if those neighbors are assigned to the same cluster with cell *i*. At the beginning connectivity value is equal 0 and increase with value 1/*i* when the *i*-th nearest neighbors is not assigned to the same cluster with cell *i*. This procedure is repeated for all cells in the dataset. In general, the higher the connectivity index is, the less separated the defined landmarks are. The connectivity index is computed using “connectivity” function implemented in R package clv. 2) coverage bias which estimates the read depth distribution in the two-dimensional embedding space; 3) sensitivity to identify rare populations. Through systematic benchmarking, we found that bin size in the range from 1kb to 10kb appeared to work well on the three benchmarks, we selected 5kb as the default bin width for all the analysis in this work (**Supplementary Methods** and **Supplementary Fig. 2**).

#### Matrix Binarization

We found that the vast majority of the elements in the cell-by-bin count matrix is “0”, indicating either closed chromatin or missing value. Among the non-zero elements, some has abnormally high coverage (> 200) perhaps due to the alignment errors. These items usually account for less than 0.1% of total non-zero items in the matrix. Thus, we change the top 0.1% elements in the matrix to “0” to eliminate potential alignment errors. We next convert the remaining non-zero elements to “1”.

#### Bin Filtering

We next filter out any bins overlapping with the ENCODE blacklist downloaded from http://mitra.stanford.edu/kundaje/akundaje/release/blacklists/. Second, we remove reads mapped to the X/Y chromosomes and mitochondrial DNA. We sort the bins based on the coverage and filter out the top 5% to remove the invariant features. Please note that we do not perform coverage-based bin filtering for a dataset that has low coverage (average fragment number less than 5,000) where the ranking of bin may be fluctuated by the noise.

#### Dimensionality Reduction

We next apply the following dimensionality reduction method to project the high-dimension data to a low-dimension manifold for clustering and visualization. Now, let us express the algorithm in matrix notation. Let 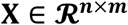 be a dataset with ***n*** cells and ***m*** bins and ***X*** = {**0,1**}. The first step is to compute a similarity matrix between the ***m*** highdimensional data points to construct the ***n***-by-***n*** pairwise similarity matrix using a kernel function ***k*** that is an appropriate similarity metric. A popular choice is gaussian kernel:

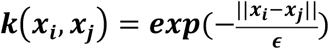

where ‖.‖ is a square root of Euclidean distance between observations ***i*** and ***j***.

Due the binarization nature of single cell ATAC-seq dataset, in this case, we replace the Gaussian kernel with Jaccard coefficient which estimates the similarity between cells simply based on ratio of overlap over the total union:

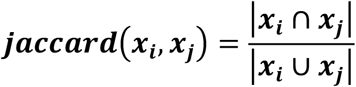

For instance, given two cells ***x_i_*** = {**0,1,1,0**} and ***x_j_*** = {**1,0,1,1**}, the Jaccard coefficient is ***jaccard***(***x_i_, x_j_***) = **1/4**. The Jaccard coefficient has the following properties that meet the requirement of being a kernel function:

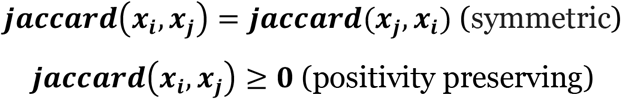

Using ***jaccard*** as a kernel function, we next form a symmetric kernel matrix 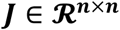 where each entry is obtained as ***J_i,j_*** = ***jaccard***(***x_i_, x_j_***)

Theoretically, the similarity ***J_i,j_*** would reflect the true similarity between cell ***x_i_*** and ***x_j_***. Unfortunately, due to the high-dropout rate, this is not the case. If there is a high sequencing depth for cell ***x_i_*** or ***x_j_***, then ***J_i,j_*** tend to have higher values, regardless whether cell ***x_i_*** and ***x_j_*** is actually similar or not.

This can be proved theoretically. Given 2 cells ***x_i_*** and ***x_j_*** and corresponding coverage (number of “1”s) 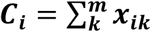 and 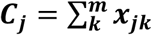, let ***P_i_ = C_i_/m***. and ***P_j_ = C_j_/m*** be the probability of observing a signal in cell ***x_i_*** and ***x_j_*** where ***m*** is the length of the vector. Assuming ***x_i_*** and ***x_j_*** are two “random” cells without any biological relevance, in another word, the “1”s in ***x_i_*** and ***x_j_*** are randomly distributed, then the expected Jaccard index between cell ***x_i_*** and ***x_j_*** can be calculated simply as:

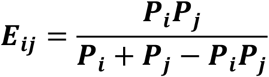

Because ***P_i_ × P_j_* > 0** (no empty cells allowed), then

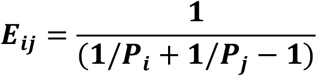

The increase of either ***P_i_*** or ***P_j_*** will result in an increase of ***E_ij_*** which suggests the Jaccard similarity between cells is highly affected by the read depth. Such observation prompts us to develop an *ad hoc* normalization method to eliminate the read depth effect.

To learn the relationship between the ***E_ij_*** and ***J_ij_*** from the data, we next fit a curve to predict the observed Jaccard coefficient ***J_ij_*** as a function of its expected value ***E_ij_*** by fitting a polynomials regression of degree 2 using R function lm. Theoretically, ***E_ij_*** should be linear with ***J_ij_*** if cells are completely random, but in real dataset, we have observed a non-linearity between ***E_ij_*** and ***J_ij_*** especially among the high-coverage cells. We suspect, to some extent, the degree of randomness of fragment distribution in a single cell is associated with the coverage. To better model the non-linearity, we include a second order polynomial in our model:

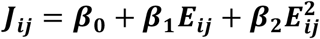

This fitting provided estimators of parameters 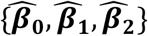. As such, we next use it to normalize the observed Jaccard coefficient by:

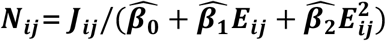

The fitting of the linear regression, however, can be very time consuming with a large matrix. Here we test the possibility of performing this step on a random subset of ***y*** cells in lieu of the full matrix. When selecting a subset of ***y*** cells to speed up the first step, we do not select cells at random with a uniform sampling probability. Instead, we set the probability of selecting a cell ***i*** to

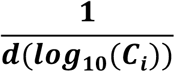

where ***d*** is the density estimate of all log10-transformed cell fragment count and ***C_i_*** is the number of fragments in cell ***i*** and 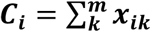. Similar approach was first introduced in SCTranscform^51^ to speed up the normalization of single cell RNA-seq.

We then proceed to normalize the full Jaccard coefficient matrix 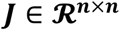 using the regression model learned from ***y*** cells and compared the results to the case where all cells are used in the initial estimation step as well. We use the correlation of normalized Jaccard coefficient to compare this partial analysis to the full analysis. We observe that using as few as 2000 cells in the estimation gave rise to virtually identical estimates. We therefore use 2,000 cells in the initial model-fitting step. To remove outliers in the normalized similarity, we use the 0.99 quantile to cap the maximum value of the normalized matrix.

Next, using normalized Jaccard coefficient matrix ***N***, we next normalize the matrix by:

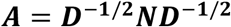

where 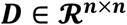 is a diagonal matrix which is composed as 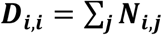. We next perform eigenvector decomposition against ***A***.

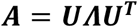

The columns 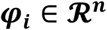 of 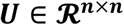 are the eigenvectors. The diagonal matrix 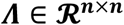 has the eigenvalues ***λ*_1_ ≥ *λ*_2_ ≥ … ≥ 0** in descending order as its entries. Finally, we report the first ***r*** eigenvectors as the final low-dimension manifold.

#### Evaluation of Ad Hoc Normalization Method

To assess the performance of normalization of SnapATAC we processed three datasets. As shown in **Supplementary Fig. 3**, before normalization, SnapATAC exhibits a strong gradient that is correlated with sequencing depth within the cluster (**Supplementary Fig. 3a**). Although the sequencing depth effect is still observed in some of the small clusters, it is clear that the normalization method has largely eliminated the read depth effect as compared to the unnormalized ones (**Supplementary Fig. 3b**).

To better quality the coverage bias, we next computed the Shannon entropy that estimates the “uniformness” of the distribution of cell coverage in the UMAP embedding space. In detail, we first chose the top 10% cells of the highest coverage as “high-coverage” cells. Second, in the 2D UMAP embedding space, we discretize “high-coverage” cells from a continuous random coordinate (umap1, umap2) into bins (n=50) and returns the corresponding vector of counts. This is done using a function called “discretize2d” in the “entropy” R package. Third, we estimated the Shannon entropy of the random variable from the corresponding observed counts. This is done using function “entropy” in the “entropy” R package. A higher entropy indicates that the “high-coverage” cells are more uniformly distributed in the UMAP embedding space, overall suggesting a better normalization performance.

We next examine another eight possible sources of biases by projecting to the UMAP embedding space, some metrics show cluster specificity for all three methods perhaps due to biological relevance, but all three methods can reveal significant biological heterogeneity without exhibiting substantial intra-cluster bias for any metrics examined (**Supplementary Fig. 4**).

#### Removing batch effects using Harmony

When the technical variability is at a larger scale than the biological variability, we apply batch effect corrector - Harmony^23^ - to eliminate such confounding factor. Given two datasets **X = {X^1^, X^2^}** generated using different technologies, we first calculate the joint low-dimension manifold ***U* = {*U*^1^, *U*^2^}** as described above. We next apply Harmony to ***U*** to regress out batch effect, resulting in a new harmonized embedding ***U^H^***. This is implemented as a function “runHarmony” in SnapATAC package.

#### Selection of Eigenvector and Eigenvalues

We next determine how many eigenvectors to include for the downstream analysis. Here we use an *ad hoc* approach for choosing the optimal number of components. We look at the scatter plot between every two pairs of eigenvectors and choose the number of eigenvectors that start exhibiting “blob”-like structure in which no obvious biological structure is revealed.

#### Nyström Landmark-Extension

The computational cost of the dimensionality reduction scales quadratically with the increase of number of cells. For instance, calculating and normalizing the pair-wise kernel Matrix *N* becomes computationally infeasible for large-scale dataset. To overcome this limitation, here we combine the Nyström method^19,52^ (a sampling technique) and our dimensionality reduction method to present Nyström landmark-extension method.

A Nyström landmark-extension algorithm includes three major steps: i) sampling ***O***(***K***): sample a subset of ***K*(*K ≪ N*)** cells from ***N*** total cells as “landmarks”. Instead of random sampling, here we adopt a density-based sampling approach developed in SCTransform^51^ to preserve the density distribution of the ***N*** original points; ii) embedding ***O*(*K*^2^)**: compute the low-dimension embedding for ***K*** landmarks; iii) extension ***O*(*N − K*)**: project the remaining ***N − K*** cells onto the low-dimensional embedding as learned from the landmarks to create a joint embedding space for all cells.

This approach significantly reduces the computational complexity and memory usage given that ***K*** is considerably smaller than ***N***. The out-of-sample extension (step iii) further enables projection of new single cell ATAC-seq datasets to the existing reference single cell atlas. This allows us to further develop a supervised approach to predict cell types of a new single cell ATAC-seq dataset based on an existing reference atlas.

A key aspect of this method is the procedure according to which cells are sampled as landmark cells, because different sampled landmark cells give different approximations of the original embedding using full matrix. Here we employ the density-based sampling as described above which preserves the density distribution of the original points.

Let 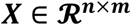 be a dataset with ***n*** cells and ***m*** variables (bins) and 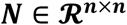 be a symmetric kernel matrix calculated using normalized Jaccard coefficient. To avoid calculating the pairwise kernel matrix and performing eigen-decomposition against a big matrix 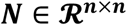, we first sample ***k*(*k ≪ n*)** landmarks without replacement. This breaks down the original kernel matrix 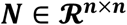 into four components.

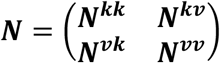

in which 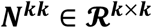 is the pairwise kernel matrix between ***k*** landmarks and 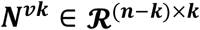 is the similarity matrix between **(*n − k*)** cells and ***k*** landmarks. Using ***N^kk^***, we perform dimensionality reduction to obtain the ***r***-rank manifold 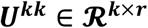 as described above.

Using ***N^vk^*** which estimates the similarity between ***n − k*** cells and ***k*** landmark cells, we project the rest of ***n − k*** cells to the embedding previously obtained using ***k*** landmark:

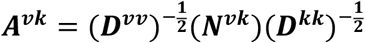

where 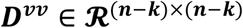 is a diagonal matrix which is composed as 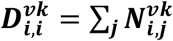. The projected coordinates of the new points onto the r-dimensional intrinsic manifold defined by the landmarks are then given by,

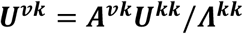

The resulting 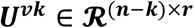 is the approximate ***r***-rank low dimension representation of the rest ***n − k*** cells. Combing ***U^kk^*** and ***U^vk^***creates a joint embedding space for all cells:

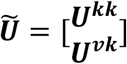

In the approximate joint ***r***-rank embedding space 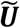, we next create a k-nearest neighbor (KNN) graph in which every cell is represented as a node and edges are drawn between cells within k nearest neighbors defined using Euclidean distance. Finally, we apply community finding algorithm such as Louvain (implemented by igraph package in R) to identify the ‘communities’ in the resulting graph which represents groups of cells sharing similar profiles, potentially originating from the same cell type.

#### Optimizing the Number of Landmarks

To evaluate the effect of the number of landmarks, we apply our method to a complex dataset that contains over 80k cells from 13 different mouse tissues. We employ the following three metrics to evaluate the performance. First, using different number of landmarks (k) ranging from 1,000 to 10,000, we compare the clustering outcome to the cell type label defined in the original study. The goal of this is to identify the “elbow” point that performance drops abruptly. Second, for each sampling, we repeat for five times using different set of landmarks to evaluate stability between sampling. Third, we spiked in 1% Patski cells to assess the sensitivity of identifying rare cell types. We choose Patski cells because these cells were profiled using the same protocol by the same group (Data source listed in **Supplementary Table S1**) to minimize the batch effect.

We observe that using as few as 5,000 landmarks can largely recapitulate the result obtained using 10,000 landmarks (**Supplementary Fig. 5a**), and 10,000 landmarks can achieve highly robust embedding between sampling (**Supplementary Fig. 5b**) and successfully recover spiked-in rare populations (**Supplementary Fig. 5c**). To obtain a reliable low-dimensional embedding, we use 10,000 landmarks for all the analysis performed in this study.

#### Ensemble Nyström Method

Nyström method is stochastic in its nature, different sampling will result in different embedding and clustering outcome. To improve the robustness of the clustering method, we next employ Ensemble Nyström Algorithm which combines a mixture of Nyström approximation to create an ensemble representation^53^. Supported by theoretical analysis, this Ensemble approach has been demonstrated to guarantee a convergence and in a faster rate in comparison to standard Nyström method^53^. Moreover, this ensemble algorithm naturally fits within distributed computing environments, where their computational costs are roughly the same as that of the standard Nyström single sampling method.

We treat each approximation generated by the Nyström method using ***k*** landmarks as an expert and combined ***p* ≥ 1** such experts to derive an improved approximation, typically more accurate than any of the original experts^53^.

The ensemble set-up is defined as follows. Given a dataset 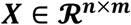 of ***n*** cells. Each expert ***S_j_*** receives ***k*** landmarks randomly selected from matrix ***X*** using density-based sampling approach without replacement. Each expert ***S_r_, r* ∈ [1, *p*]** is then used to define the low dimension embedding 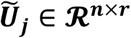 as described above. For each low-dimension embedding 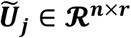, we create a KNN-graph as 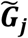. Thus, the general form of the approximation, 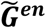, generated by the ensemble Nyström method is

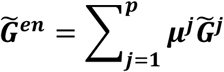

where ***μ^j^*** is the mixture weights that can be defined in many ways. Here we choose to use the most straightforward method by assigning an equal weight to each of the KNN-graph obtained from different samplings, ***μ^j^* = 1/*p, r* ∈ [l, *p*]**. While this choice ignores the relative quality of each Nyström approximation, it is computational efficient and already generates a solution superior to any one of the approximations used in the combination. Using the ensemble weighted KNN graph 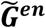, we next apply community finding algorithm to identify cell clusters. By testing on the mouse atlas dataset^8^, we demonstrate that the clustering stability of the ensemble approach is significantly higher than the standard Nystrom method (**Supplementary Fig. 5d**).

#### Visualization

We use the t-SNE implemented by FI-tsne, Rtsne or UMAP (umap_0.2.0.0) to visualize and explore the dataset.

#### Gene Accessibility Score

To annotate the identified clusters, SnapATAC calculated the gene-body accessibility matrix ***G*** using “calGmatFromMat” function in SnapATAC packge where ***G_ij_*** is the number of fragments overlapping with **j**-th genes in **i**-th cell. ***G_ij_*** is then normalized to CPM (count-per-million reads) as 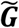. The normalized accessibility score is then smoothed using Markov affinity-graph based method:

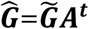

where ***A*** is the adjacent matrix obtained from K nearest neighbor graph and ***t*** is number of steps taken for Markov diffusion process. We set ***t* = 3** in this study. Please note that the gene accessibility score is only used to guide the annotation of cell clusters identified using cell-by-bin matrix. The clusters are identified using cell-by-bin matrix in prior.

#### Read Aggregation & Peak Calling

After annotation, cells from the same cluster are pooled to create aggregated signal for each of the identified cell types. This allows for identifying *cis*-elements from each cluster. MACS2 (version 2.1.2) is used for generating signal tracks and peak calling with the following parameters: --nomodel --shift 100 --ext 200 --qval 1e-2 -B –SPMR. This can be done by “runMACS” function in SnapATAC package.

#### Motif Analysis

SnapATAC incorporates chromVAR^14^ to estimate the motif variability and Homer^21^ for *de novo* motif discovery. This is implemented as function “runChromVAR” and “runHomer” in SnapATAC package.

#### Identification of differentially accessible peaks

For a given group of cells *C_i_*, we first look for their neighboring cells *C_j_* (|*C_i_*| = |*C_j_*|) in the low-dimension manifold as “background” cells to compare to. If *C_i_* accounts for more than half of the total cells, we use the remaining cells as local background. Next, we aggregate *C_i_* and ćjo to create two raw-count vectors as *V_ci_* and *V_c_j*. We then perform differential analysis between *V_ci_* and *V_cj_* using exact test as implemented in R package edgeR (v3.18.1) with BCV=0.1. P-value is then adjusted into False Discovery Rate (FDR) using Benjamini-Hochberg correction. Peaks with FDR less than 0.01 are selected as significant DARs. However, the statically significance is under powered for small clusters.

#### GREAT analysis

SnapATAC incorporates GREAT analysis^39^ to infer the candidate biological pathway active in each cell populations. This is implemented as function “runGREAT” SnapATAC package.

#### Integration with single cell RNA-seq

We use canonical correlation analysis (CCA) embedded in Seurat V3^18^ to integrate single cell RNA-seq and single cell ATAC-seq. We first calculate the gene accessibility account at variable genes identified using single cell RNA-seq dataset. This can be done using a function called “createGmatFromMat” in SnapATAC package. Next, SnapATAC converts the snap object to a Seurat v3 object using a function called “SnapToSeurat” in preparation for integration. Different from integration method in Seurat, we use the our low-dimension manifold as the dimensionality reduction method in the Seurat object. We next follow the vignette in Seurat website (https://satijalab.org/seurat/v3.0/atacseq_integration_vignette.html) to integrate these two modalities. The cell type for scATAC-seq is predicted using function “TransferData” in Seurat V3.

Finally, for each single cell ATAC profile, we infer its gene expression profile by calculating the weighted average expression profile of its nearest neighboring cells in the single cell RNA-seq dataset^18^. By doing so, we create pseudo-cells that contain information of both chromatin accessibility and gene expression profiles. The imputation of gene expression profile is done by “TransferData” function in Seurat V3.

#### Linking enhancers to putative target genes

Using the “pseudo” cells, we next sought to predict the putative target genes for regulatory elements based on the association between expression of a gene and chromatin accessibility at its enhancer elements. Given a gene ***G***, we first identify its surrounding regulatory elements within 1MB window flanking ***G***. Let ***Y^G^*** be the imputed gene expression value for gene ***G*** among ***n*** cells. We perform logistic regression using ***Y^G^*** as variable to predict the binary state for each of peaks surrounding ***G***. The idea behind using logistic regression is that if there is a relationship between the gene expression (continuous variable) and chromatin accessibility (categorical variable), we should be able to predict chromatin accessibility from the gene expression. Logistic regression does not make many of the key assumptions such as normality of the continuous variables. In addition, since we only have one variable (gene expression) for prediction every time, there is no problem of multicollinearity.

We next fit logistic regression between each of flanking peak and gene expression using “glm” function in R with binomial(link=‘logit’) as the family function. By doing so, we obtain the regression coefficient ***β*_1_** and its corresponding P-value for each peak separately. Here we used 5e-8, a standard P-value cutoff for human genome-wise association study to determine the significant association. While this cutoff is less sample or gene specific compared to more complicated methods such as permutation test, it is computational efficient and already generates a reasonable set of gene-enhancer pairings.

To evaluate the performance of our methods, we compare our prediction with cis-eQTL derived from interferon-γ and lipopolysaccharide stimulation of monocytes^25^. Significant cis-eQTL associations are downloaded from supplementary material (Table S2) in Fairfax (2014)^25^. We filter cis-eQTL based on two criteria: 1) only cis-eQTLs that overlap with the peaks identified in PBMC dataset are considered; 2) In addition, we only keep the cis-eQTLs whose genes overlap with the variable genes determined by scRNA-seq. This filtering reduced the cis-eQTL list to 456 candidates.

Next, we estimate the association for each of cis-eQTLs by preforming logistic regression test as described above. To make a comparison, we derive a set of negative pairs matched for the distance. The negative control pairs for cis-eQTL are chosen in the following manner to control for both distance and chromatin accessibility: for each positive eQTL pair *p_ij_* which connects gene *i* and enhancer *j* with a distance of *d_ij_*, we look for the enhancer *k* on the opposite direction of the gene *i* that minimizes |*d_ij_ − d_iz_*|. By doing so, the negative sets are controlled for distance, chromatin accessibility level and gene expression level.

### Simulation of scATAC-seq datasets

First, we download the alignment files (bam files) for ten bulk ATAC-seq experiment from ENCODE (data source listed in **Supplementary Table S2**). From each bam file, we simulate 1,000 single cell ATAC-seq datasets by randomly down sampling to a variety of coverages ranging from 1,000 to 10,000 reads per cells. We next create a cell-by-bin matrix of 5kb which is used for SnapATAC clustering. Merging peaks identified from each bulk experiment, we create cell-by-peak matrix used for LSA, Cis-Topic, Cicero and chromVAR for clustering. We repeat the sampling for n=10 times to estimate the variability of the clustering.

### Comparison of scalability

To compare the scalability between SnapATAC to other methods, we next simulate multiple datasets of different number of cells ranging from 20k to 1M. We simulate these datasets in the following manner. Using the 80k mouse atlas dataset, we randomly sample this dataset to different number of cells ranging from 20k t0 1M cells. For the sampling that has cells more than 80K, we sample with replacement and introduce perturbation to each cell by randomly removing 1% of the “1”s in each of the cells. This removes the duplicate cells and largely maintains the density of the matrix.

For each sampling, we then perform dimensionality reduction using LSA and cisTopic and compare their CPU running time. Specifically, we monitor the running time for 1) TF-IDF transformation and Singular Value Decomposition (SVD) for LSA, 2) function “runModels” with topics = c(2, 5, 10, 15, 20, 25, 30, 35, 40) and “selectModel” function in cisTopic. The time for matrix loading is not counted.

All the comparisons were tested on a machine with 5 AMD Operon (TM) Processor 6276 CPUs.

### Doublets Detection Using Scrublet

To identify doublets from secondary motor cortex single nucleus ATAC-seq datasets, we use single cell RNA-seq doublets detection algorithm Scrublet^37^. Briefly, Scrublet identifies doublets in the following manner: 1) Scrublet performs normalization, gene filtering, and principal components analysis (PCA) to project the high-dimension data to a low-dimension space; 2) Scrublet simulates doublets by adding the unnormalized counts from randomly sampled observed transcriptomes; 3) the simulated doublets are projected to the low dimension embedding computed in step 1. The more neighbors of a cell are the simulated doublets, the more likely this cell is a “doublet”. Based on this idea, a KNN classifier was then used to estimate the doublet score for each cell.

Since Scrublet was designed for detecting doublets in single cell RNA-seq, it is unclear whether it can be used for single cell ATAC-seq. To examine this, we applied Scrublet to a single cell ATAC-seq dataset of mixed human and mouse cells where the “ground-truth” doublets can be identified based on the alignment ratio to human and mouse genome. Compared to the ground truth, Scrubet can identify over 90% of the doublets in this dataset with ∼90% accuracy (**Supplementary Fig. 26**). This result suggests that although Scrubet was not developed for detecting doublets in single cell ATAC-seq, it can find the doublets in scATAC-seq dataset with reasonable accuracy and sensitivity.

### Projection of single cell ATAC-seq datasets to reference atlas

We reason that landmark-extension algorithm can also be extended to project new single cell ATAC-seq datasets to a reference atlas. Given a query dataset 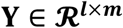 that contains ***l*** query cells with ***m*** bins and a reference dataset 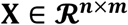 with ***n*** reference cells of ***m*** bins. We first randomly sample ***k*** =10,000 landmarks from **X** using density-based sampling as described above. Next, we compute the pairwise similarity using normalized jaccard coefficient for ***k*** landmarks as 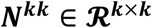 and obtain the low-dimension manifold 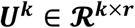. We then compute 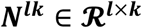 which estimates the similarity between ***l*** query cells and ***k*** landmark cells, and then project the ***l*** query cells to the embedding pre-computed for ***k*** landmark cells as following:

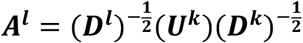

where 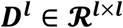 is a diagonal matrix which is composed as 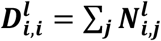 and 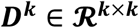 is a diagonal matrix which is composed as 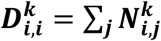

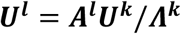

The resulting 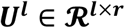 is the predicted low-dimension manifold for ***l*** query cells.

In the joint embedding space **[*U^k^,U^l^*]**, we next identify the mutual nearest neighbors between query and landmark cells. For each cell *i*_1_ ∈ **X^*k*^** belonging to the landmarks, we find the ***k. nearest*(5)** cells in the query dataset with the smallest distances to ***i*_1_**. We do the same for each cell in query cell dataset to find its ***k. nearest*** (5) neighbors in the landmark dataset. If a pair of cells from each dataset is contained in each other’s nearest neighbors, those cells are considered to be mutual nearest neighbors or MNN pairs (or “anchors”). We interpret these pairs as containing cells that belong to the same cell type or state despite being generated in both landmark and query cells. Thus, any differences between cells in MNN pairs should theoretically represent the non-overlapping cell types. Here we removed any query cells that failed to identify an MNN pair correspondence in the reference dataset.

To make a classification of the remaining query cells according to the reference dataset, we next apply the neighborhood-based classifier and wish to highlight the pioneering work by Seurat V3^18^. First, we score each anchor (or MNN pair) using shared nearest neighbor (SNN) graph by examining the consistency of edges between cells in the same local neighborhood as described in the original study^18^. Second, we define a weight matrix that estimates the strength of association between each query cell ***c***, and each landmark ***i***. For each query cell ***c***, we identify the nearest ***s*** landmarks in the reference dataset in the joint embedding space. Nearest anchors are then weighted based on their distance to the cell ***c*** over the distance to the ***s***-th anchor cell. For each cell ***c*** and anchor ***i***, we compute the weighted similarities as:

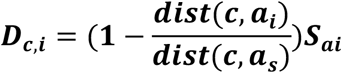

Where ***dist*(*c, i*)** is the Euclidean distance in the joint embedding space and *S_ai_* is the weight for the corresponding MNN pair (anchor). We then normalize the similarity using exponential function:

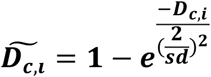

where sd is set to 1 by default. Finally, we normalize across all ***s*** anchors:

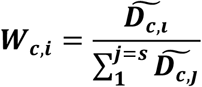

Here we set ***s = 50***. Please note that the similarity to cells beyond the ***s^th^*** anchor neighbor is set to be zero.

Let 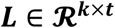 be the binary label matrix for ***k*** landmarks with ***t*** clusters. ***L_ij_* = 1** indicates the class label for ***i***-th landmark cell is ***j***-th cluster. The row sum of ***L*** must be 1, suggesting each landmark cell can only be assigned to one cluster label. We then compute label predictions for query cells as ***P^l^***:

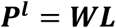

The resulting ***P^l^*** is a probability matrix within 0 and 1, 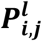 indicates the probability of a cell ***i*** belong to ***j*** cluster. Similarly, we infer the t-SNE position of query cells by replacing ***L*** with t-SNE coordinates of reference points. It is important to note that the distance between cells in the inferred t-SNE coordinate does not neccessarily reflect the cell-to-cell relationship.

### Tissue collection & nuclei isolation

Adult C57BL/6J male mice were purchased from Jackson Laboratories. Brains were extracted from P56-63 old mice and immediately sectioned into 0.6 mm coronal sections, starting at the frontal pole, in ice-cold dissection media. The secondary motor cortex (MOs) region was dissected from the first three slices along the anterior-posterior axis according to the Allen Brain reference Atlas (http://mouse.brain-map.org/, see **Supplementary Fig. 15a** for depiction of posterior view of each coronal slice; dashed line highlights the MOs regions on each slice). Slices were kept in ice-cold dissection media during dissection and immediately frozen in dry ice for posterior pooling and nuclei production. For nuclei isolation, the MOs dissected regions from 15-23 animals were pooled, and two biological replicates were processed for each slice. Nuclei were isolated as described in previous studies^54,55^, except no sucrose gradient purification was performed. Flow cytometry analysis of brain nuclei was performed as described in Luo et al^54^.

### Tn5 transposase purification & loading

Tn5 transposase was expressed as an intein chitin-binding domain fusion and purified using an improved version of the method first described by Picelli et al^56^. T7 Express lysY/I (C3013I, NEB) cells were transformed with the plasmid pTXB1-ecTn5 E54K L372P (#60240, Addgene)^56^. An LB Ampicillin culture was inoculated with three colonies and grown overnight at 37°C. The starter culture was diluted to an OD of 0.02 with fresh media and shaken at 37°C until it reached an OD of 0.9. The culture was then immediately chilled on ice to 10°C and expression was induced by adding 250 μM IPTG (Dioxane Free, CI8280-13, Denville Scientific). The culture was shaken for 4 hours at 23°C after which cells were harvested in 2 L batches by centrifugation, flash frozen in liquid nitrogen and stored at −80°C. Cell pellets were resuspended in 20 ml of ice cold lysis buffer (20 mM HEPES 7.2-KOH, 0.8 M NaCl, 1 mM EDTA, 10% Glycerol, 0.2% Triton X-100) with protease inhibitors (Complete, EDTA-free Protease Inhibitor Cocktail Tablets, 11873580001, Roche Diagnostics) and passed three times through a Microfluidizer (lining covered with ice water, Model 110L, Microfluidics) with a 5 minute cool down interval in between each pass. Any remaining sample was purged from the Microfluidizer with an additional 25 ml of ice-cold lysis buffer with protease inhibitors (total lysate volume ∼50ml). Samples were spun down for 20 min in an ultracentrifuge at 40K rpm (L-80XP, 45 Ti Rotor, Beckman Coulter) at 4°C. ∼45 ml of supernatant was combined with 115 ml ice cold lysis buffer with protease inhibitors in a cold beaker (total volume = 160 ml) and stirred at 4°C. 4.2ml of 10% neutralized polyethyleneimine-HCl (pH 7.0) was then added dropwise. Samples were spun down again for 20 min in an ultracentrifuge at 40K rpm (L-80XP, 45 Ti Rotor, Beckman Coulter) at 4°C. The pooled supernatant was loaded onto ∼10ml of fresh Chitin resin (S6651L, NEB) in a chromatography column (Econo-Column (1.5 × 15 cm), Flow Adapter: 7380015, Bio-Rad). The column was then washed with 50-100 ml lysis buffer. Cleavage of the fusion protein was initiated by flowing ∼20ml of freshly made elution buffer (20 mM HEPES 7.2-KOH, 0.5 M NaCl, 1 mM EDTA, 10% glycerol, 0.02% Triton X-100, 100mM DTT) onto the column at a speed of 0.8ml/min for 25 min. After the column was incubated for 63 hrs at 4°C, the protein was recovered from the initial elution volume and a subsequent 30 ml wash with elution buffer. Protein-containing fractions were pooled and diluted 1:1 with buffer [20 mM HEPES 7.2-KOH,1 mM EDTA, 10% glycerol, 0.5mM TCEP) to reduce the NaCl concentration to 250mM. For cation exchange, the sample was loaded onto a 1ml column HiTrap S HP (17115101, GE), washed with Buffer A (10mM Tris 7.5, 280 mM NaCl, 10% glycerol, 0.5mM TCEP) and then eluted using a gradient formed using Buffer A and Buffer B (10mM Tris 7.5, 1M NaCl, 10% glycerol, 0.5mM TCEP) (0% Buffer B over 5 column volumes, 0-100% Buffer B over 50 column volumes, 100% Buffer B over 10 column volumes). Next, the protein-containing fractions were combined, concentrated via ultrafiltration to ∼1.5 mg/mL and further purified via gel filtration (HiLoad 16/600 Superdex 75 pg column (28989333, GE)) in Buffer GF (100mM HEPES-KOH at pH 7.2, 0.5 M NaCl, 0.2 mM EDTA, 2mM DTT, 20% glycerol). The purest Tn5 transposase-containing fractions were pooled and 1 volume 100% glycerol was added to the preparation. Tn5 transposase was stored at −20°C.

To generate Tn5 transposomes for combinatorial barcoding assisted single nuclei ATAC-seq, barcoded oligos were first annealed to pMENTs oligos (95 °C for 5 min, cooled to 14 °C at a cooling rate of 0.1 °C/s) separately. Next, 1 μl barcoded transposon (50 μM) was mixed with 7 ul Tn5 (∼7 μM). The mixture was incubated on the lab bench at room temperature for 30 min. Finally, T5 and T7 transposomes were mixed in a 1:1 ratio and diluted 1:10 with dilution buffer (50 % Glycerol, 50 mM Tris-HCl (pH=7.5), 100 mM NaCl, 0.1 mM EDTA, 0.1 % Triton X-100, 1 mM DTT). For combinatorial barcoding, we used eight different T5 transposomes and 12 distinct T7 transposomes, which eventually resulted in 96 Tn5 barcode combinations per sample^7^ (**Supplementary Table S6**).

### Bulk ATAC-seq data generation

ATAC-seq was performed on 30,000-50,000 nuclei as described previously with modifications^3^. Nuclei were thawed on ice and pelleted for 5 min at 500 x g at 4 °C. Nuclei pellets were resuspended in 30 μl tagmentation buffer (36.3 mM Tris-acetate (pH = 7.8), 72.6 mM K-acetate, 11 mM Mg-acetate, 17.6 % DMF) and counted on a hemocytometer. 30,000-50,000 nuclei were used for tagmentation and the reaction volume was adjusted to 19 μl using tagmentation buffer. After addition of 1 μl TDE1 (Illumina FC-121-1030), tagmentation was performed at 37°C for 60 min with shaking (500 rpm). Tagmented DNA was purified using MinElute columns (Qiagen), PCR-amplified for 8 cycles with NEBNext^®^ High-Fidelity 2X PCR Master Mix (NEB, 72°C 5 min, 98°C 30 s, [98°C 10 s, 63°C 30 s, 72°C 60 s] x 8 cycles, 12°C held). Amplified libraries were purified using MinElute columns (Qiagen) and SPRI Beads (Beckmann Coulter). Sequencing was carried out on a NextSeq500 using a 150-cycle kit (75 bp PE, Illumina).

### Bulk ATAC-seq data analysis

ATAC-seq reads were mapped to reference genome mm10 using BWA and *samtools* version 1.2 to eliminate PCR duplicates and mitochondrial reads. The paired-end read ends were converted to fragments. Using fragments, MACS2^57^ version 2.1.2 was used for generating signal tracks and peak calling with the following parameters: --nomodel --shift 100 --ext 200 --qval 1e-2 -B –SPMR. Library quality control for bulk ATAC-seq can be found in **Supplementary Table S7**.

### Single-nucleus ATAC-seq data generation

Combinatorial ATAC-seq was performed as described previously with modifications^5,7^. For each sample two biological replicates were processed. Nuclei were pelleted with a swinging bucket centrifuge (500 x g, 5 min, 4°C; 5920R, Eppendorf). Nuclei pellets were resuspended in 1 ml nuclei permeabilization buffer (5 % BSA, 0.2 % IGEPAL-CA630, 1mM DTT and cOmpleteTM, EDTA-free protease inhibitor cocktail (Roche) in PBS) and pelleted again (500 x g, 5 min, 4°C; 5920R, Eppendorf). Nuclei were resuspended in 500 μL high salt tagmentation buffer (36.3 mM Tris-acetate (pH = 7.8), 72.6 mM potassium-acetate, 11 mM Mg-acetate, 17.6% DMF) and counted using a hemocytometer. Concentration was adjusted to 4500 nuclei/9 μl, and 4,500 nuclei were dispensed into each well of a 96-well plate. Glycerol was added to the leftover nuclei suspension for a final concentration of 25 % and nuclei were stored at −80°C. For tagmentation, 1 μL barcoded Tn5 transposomes^7,56^ (**Supplementary Table S6**) were added using a BenchSmart™ 96 (Mettler Toledo), mixed five times and incubated for 60 min at 37 °C with shaking (500 rpm). To inhibit the Tn5 reaction, 10 μL of 40 mM EDTA were added to each well with a BenchSmart™ 96 (Mettler Toledo) and the plate was incubated at 37 °C for 15 min with shaking (500 rpm). Next, 20 μL 2 x sort buffer (2 % BSA, 2 mM EDTA in PBS) were added using a BenchSmart™ 96 (Mettler Toledo). All wells were combined into a FACS tube and stained with 3 μM Draq7 (Cell Signaling). Using a SH800 (Sony), 20 nuclei were sorted per well into eight 96-well plates (total of 768 wells) containing 10.5 μL EB (25 pmol primer i7, 25 pmol primer i5, 200 ng BSA (Sigma)^7^. Preparation of sort plates and all downstream pipetting steps were performed on a Biomek i7 Automated Workstation (Beckman Coulter). After addition of 1 μL 0.2% SDS, samples were incubated at 55 °C for 7 min with shaking (500 rpm). We added 1 μL 12.5% Triton-X to each well to quench the SDS and 12.5 μL NEBNext High-Fidelity 2× PCR Master Mix (NEB). Samples were PCR-amplified (72 °C 5 min, 98 °C 30 s, (98 °C 10 s, 63 °C 30 s, 72 °C 60 s) × 12 cycles, held at 12 °C). After PCR, all wells were combined. Libraries were purified according to the MinElute PCR Purification Kit manual (Qiagen) using a vacuum manifold (QIAvac 24 plus, Qiagen) and size selection was performed with SPRI Beads (Beckmann Coulter, 0.55x and 1.5x). Libraries were purified one more time with SPRI Beads (Beckmann Coulter, 1.5x). Libraries were quantified using a Qubit fluorimeter (Life technologies) and the nucleosomal pattern was verified using a Tapestation (High Sensitivity D1000, Agilent). The library was sequenced on a HiSeq2500 sequencer (Illumina) using custom sequencing primers, 25% spike-in library and following read lengths: 50 + 43 + 40 + 50 (Read1 + Index1 + Index2 + Read2)^7^.

---

**Figure S1.**
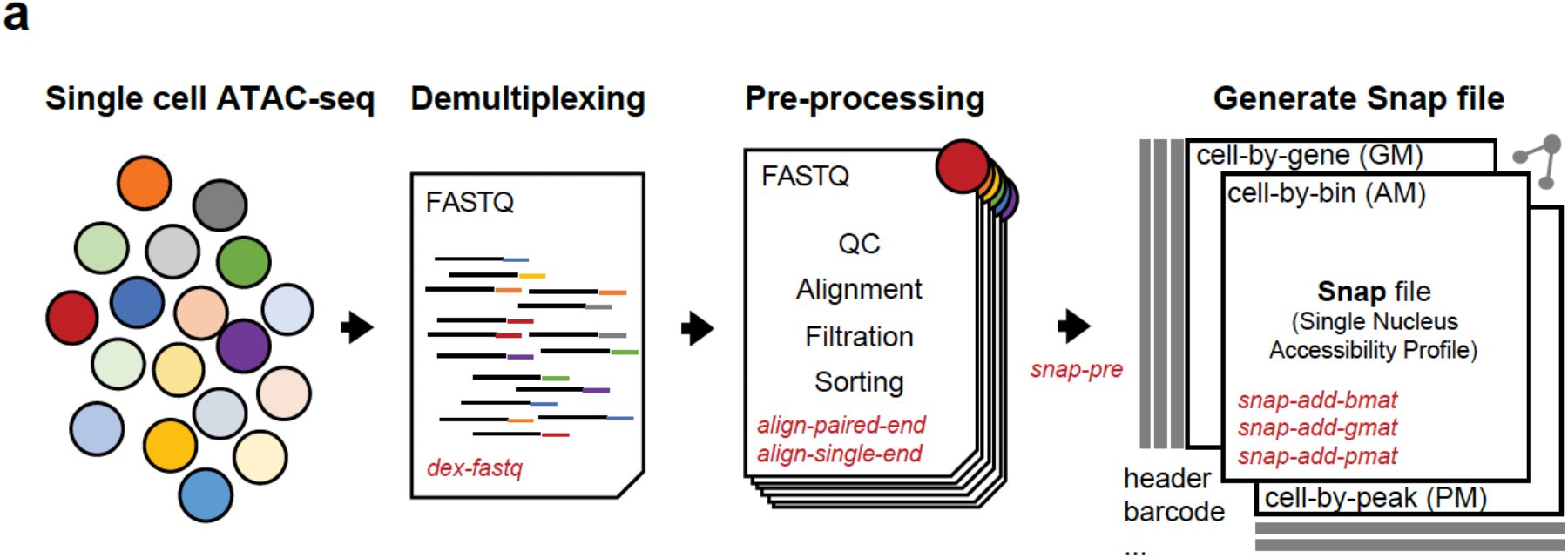
Overview of SnapTools workflow. (**a**) Demultiplexing: SnapTools first demultiplexed the fastq files by adding the cell barcodes to the beginning of each read name; Pre-processing: raw sequencing reads were aligned to the reference genome using BWA followed by filtration of erroneous alignments. A snap file was generated to store indexed reads and multiple cell matrices including cell-by-peak, cell-by-gene and cell-by-bin matrix.

**Figure S2.**
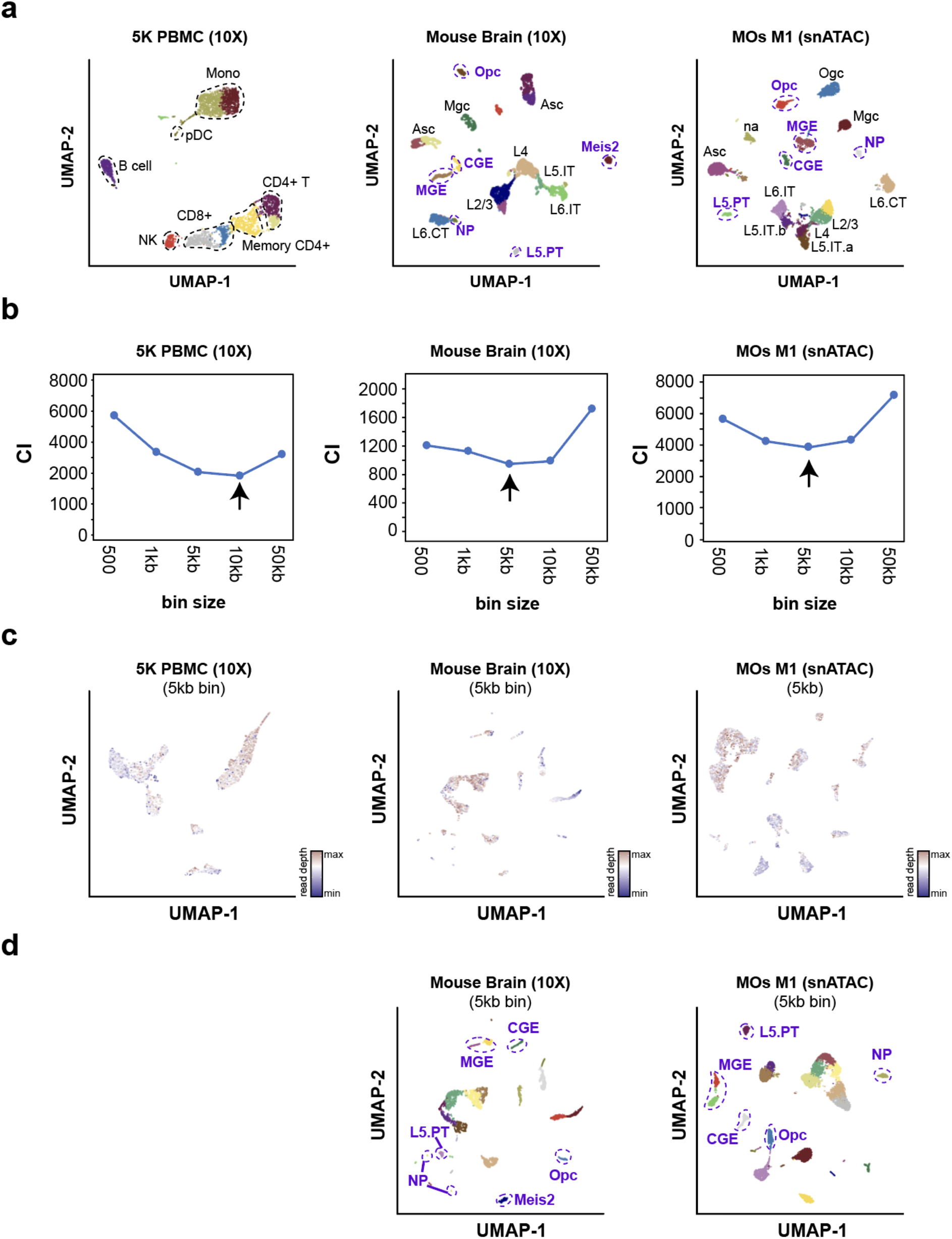
Choosing the optimal bin size. (**a**) UMAP visualization of landmark cell types identified in three benchmarking datasets. UMAP embedding was computed using cisTopic and cell types were manually annotated based on the gene accessibility score at canonical marker genes (**Supplementary Methods**). Blue dash line highlights the rare cell populations that account for less than 2% of the total population. (**b**) Relationship between connectivity index (CI) and bin sizes. Connectivity index were calculated between landmark cell types in the reduced dimension using function “connectivity” in R package “clv”. A lower CI indicates a better separation of landmark cell types. (**c**) UMAP representation of three benchmarking datasets generated using SnapATAC using 5kb bin size. Cells colored by read depth to illustrate the sequencing depth effect. (**d**) Cells are colored by cluster labels identified by SnapATAC. Data source are listed in **Supplementary Table S1**. Note that blue circles highlight rare cell populations account for less than 2% of total population.

**Figure S3.**
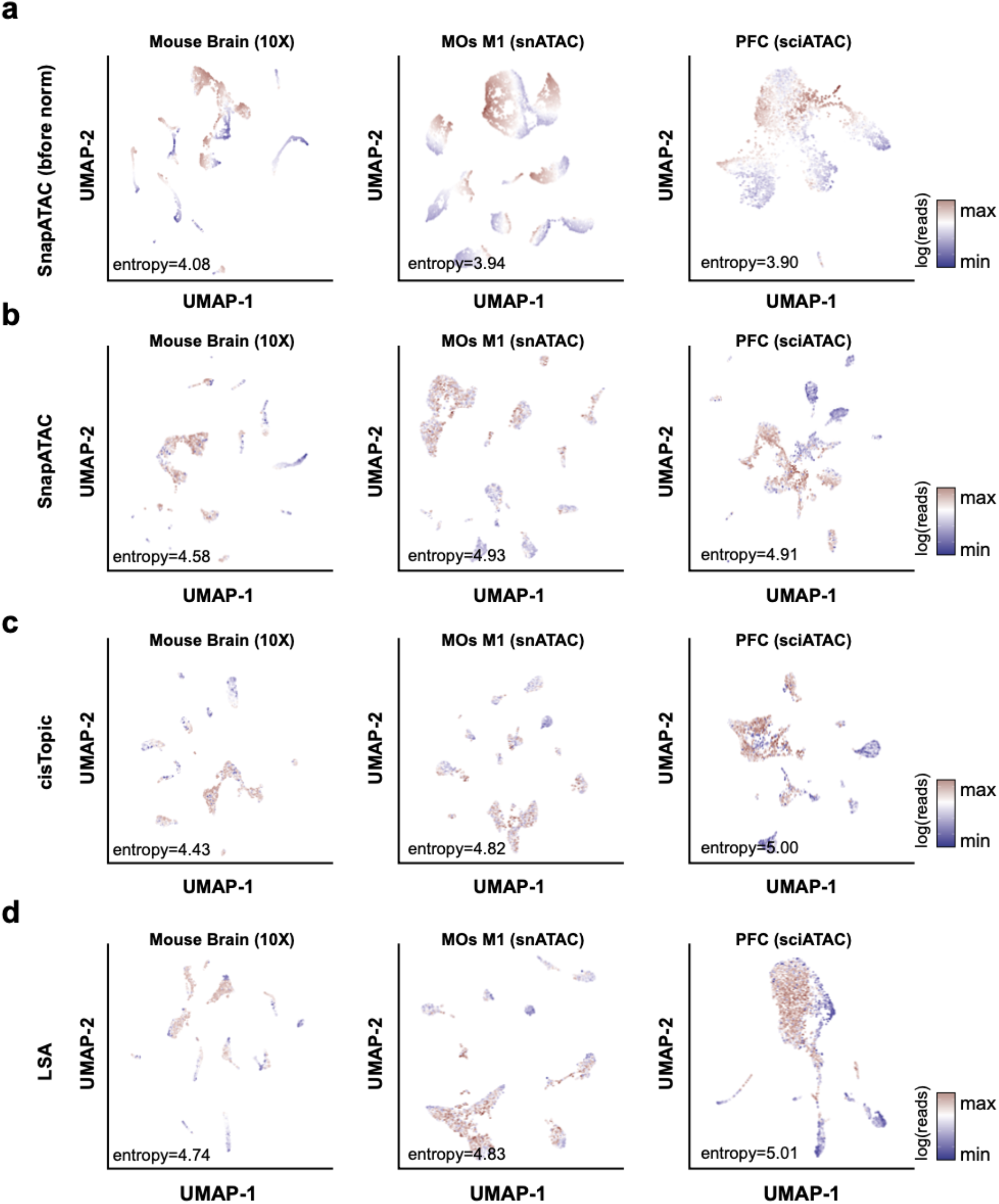
SnapATAC is robust to sequencing depth. Two dimensional UMAP representation of three benchmarking datasets analyzed by four methods (**a**) SnapATAC without normalization; (**b**) SnapATAC with normalization; (**c**) cisTopic and (**d**) Latent Sematic Analysis (LSA). Cells are color by log-scaled read depth. Read depth bias is quantified by entropy as described in the **Supplementary Methods**. Data source is listed in **Supplementary Table S1**.

**Figure S4.**
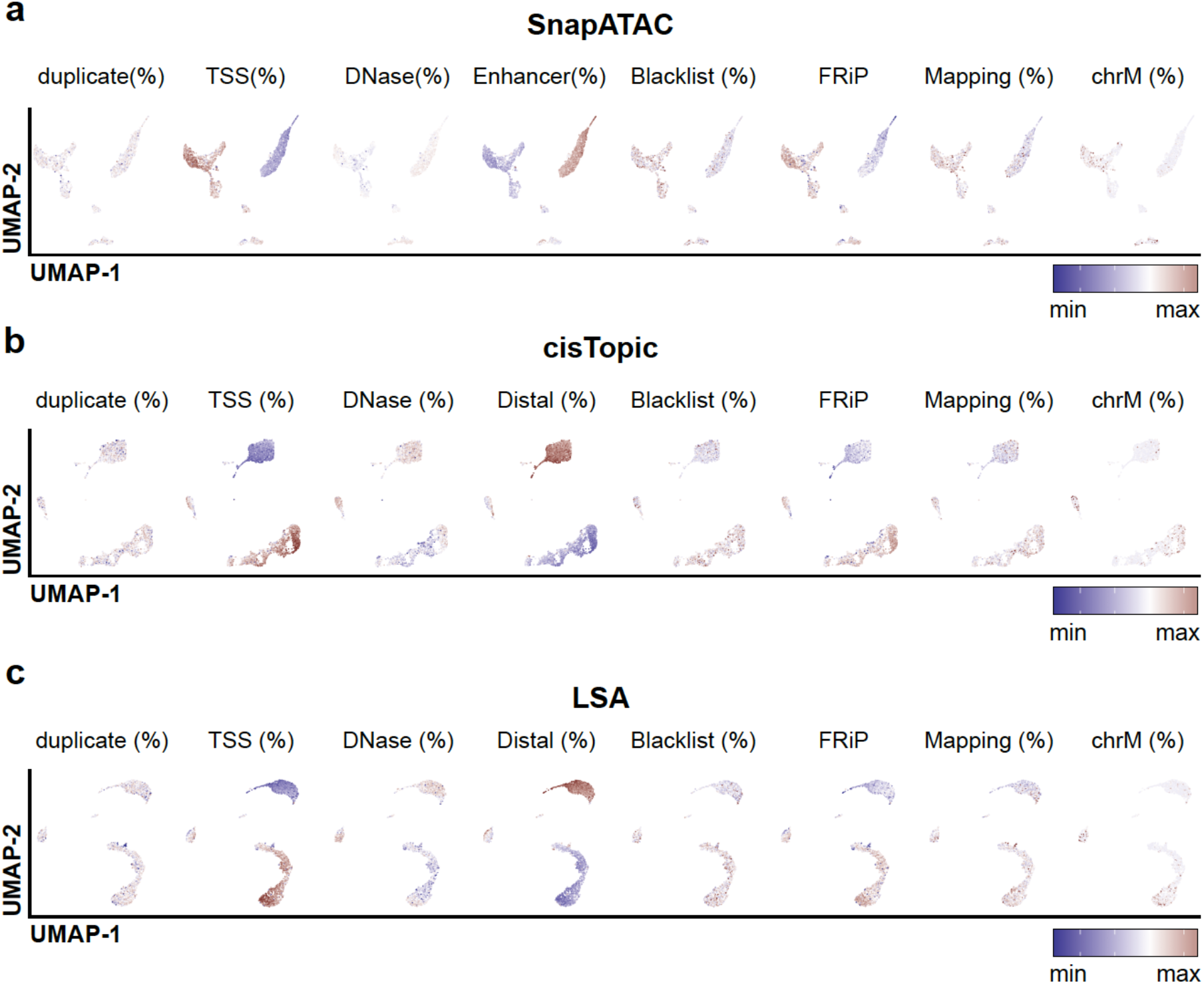
SnapATAC is robust to other biases. Potential bias in single cell ATAC-seq dataset projected onto the UMAP visualization generated using different analysis methods (**a**) SnapATAC (**b**) cisTopic and (**c**) LSA. Duplicate: percentage of fragments that are PCR duplicates. TSS: percentage of fragments overlapping or are within 1kb of a TSS. TSS position is based on the GENECODE V28 (Ensemble 92). DNase: the percentage of fragments overlapping a master DNase peak list. The DNase peak list is created by combining all ENCODE^1^ DNase peaks from hg19. Blacklist: the percentage of fragments overlapping with the ENCODE blacklist. FRiP: the percentage of fragments overlapping with the peaks defined from the aggregate signal. Mapping: the percentage of fragments that are uniquely mapped. chrM: the percentage of fragments mapped to mitochondria DNA. Dataset used in this plot is 5k PBMC (10X) as listed in **Supplementary Table S1**.

**Figure S5.**
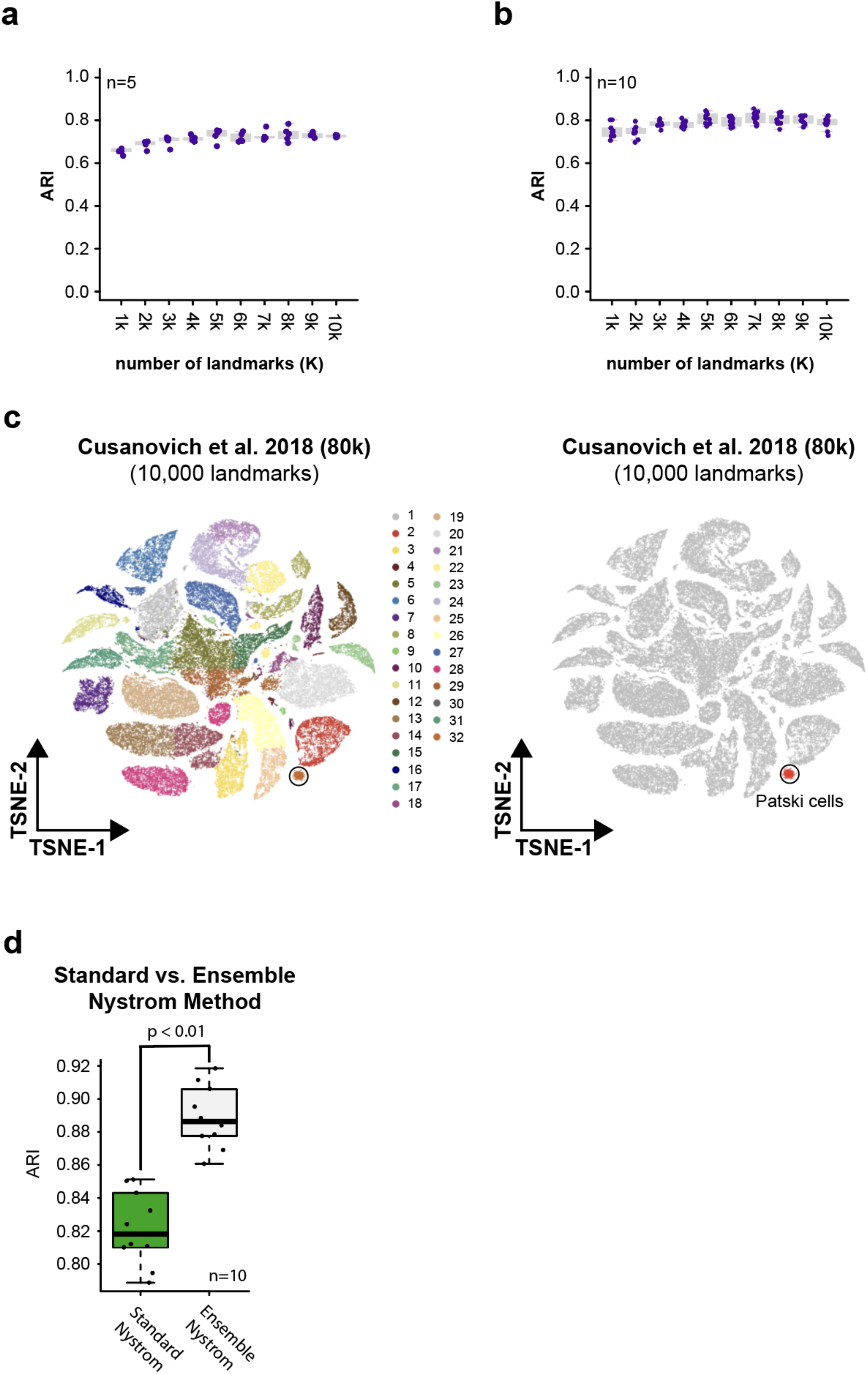
Ensemble Nyström sampling improves the scalability and stability without sacrificing the performance. (**a**) A line plot comparing the performance of clustering using various sampling parameters. The performance is evaluated using Adjusted Rank Index (ARI). SnapATAC was applied to the mouse atlas dataset that contained over 80k cells using different number of landmark cells (*k*) ranging from 1k to 10k. For each *k*, we performed clustering for n=5 times using different sets of randomly selected landmarks. (**b**) A line plot comparing the stability of clustering results between five samplings (pairwise comparison n=10). (**c**) To evaluate the sensitivity of identifying rare cell types, we spiked in 1% mouse Pastki cells generated using the same protocol in Cusanovich 2015^5^ and this rare cell population was recapitulated using 10,000 landmarks (right). (**d**) To compare the clustering reproducibility between standard and ensemble Nystrom sampling method, we performed clustering using both methods on Cusanovich 2018^8^ for five times with different randomly selected landmark cells. The clustering reproducibility quantified by ARI (adjusted rank index) between random trails is significantly higher for the ensemble Nystrom method than the standard Nystrom method (two-tailed t-test *P* < 0.01).

**Figure S6.**
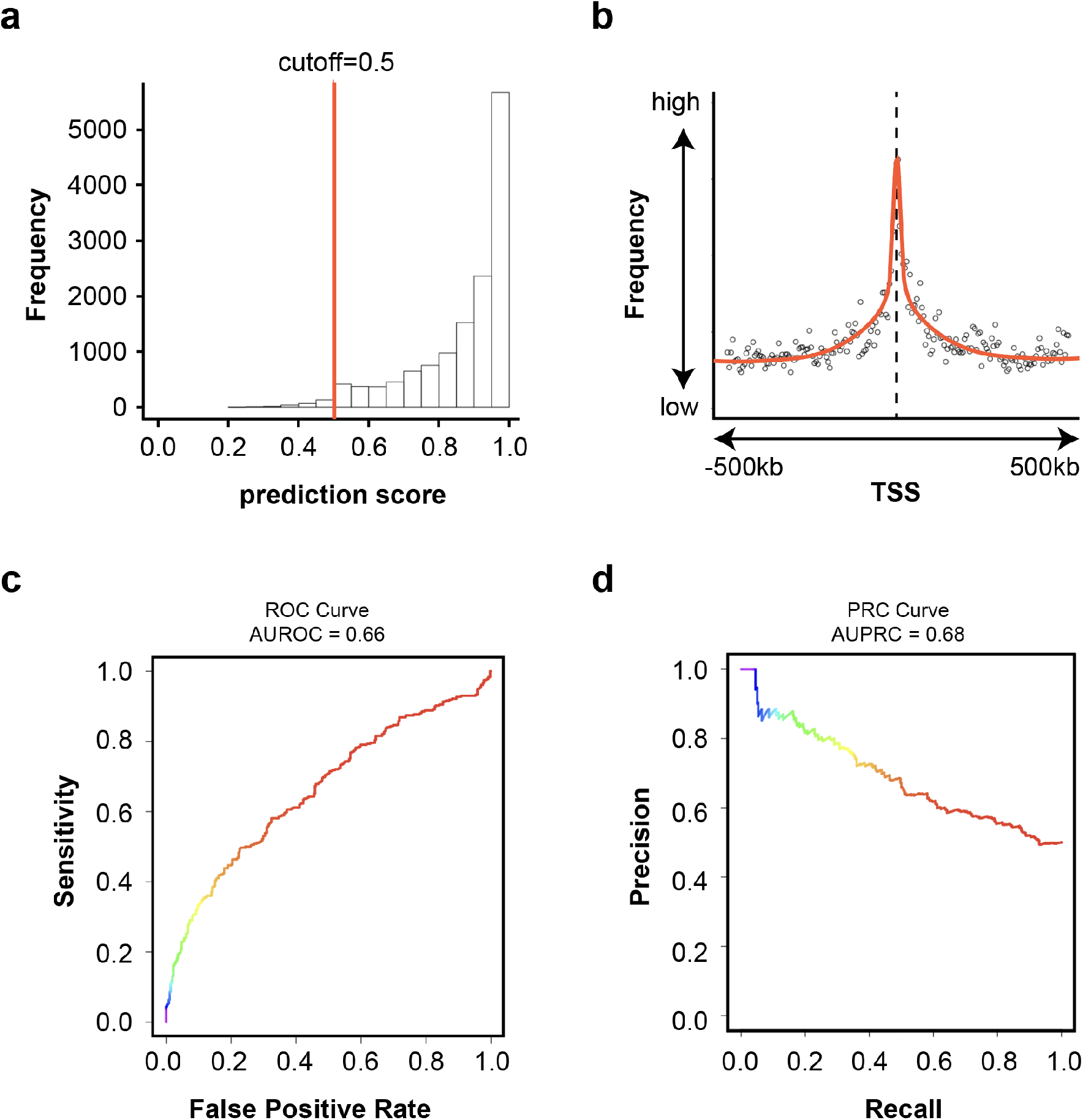
SnapATAC predicts gene and enhancer pairing by integrating scATAC-seq and scRNA-seq. (**a**) Prediction score distribution for single cell ATAC-seq (5K PBMC 10X) by SnapATAC. When predicting the cell type for scATAC-seq using corresponding scRNA-seq dataset (10X PBMC scRNA-seq), each cell in scATAC-seq was assigned with a prediction score indicating the confidence of the prediction. It ranges from 0 to 1, a higher score indicates a higher confidence. Using 0.5 as cutoff as suggested in Seurat, over 98% of cells in scATAC-seq are confidently assigned to a cell type defined in scRNA-seq. (**b**) Distance decay curve for the association (-logPvalue) between regulatory elements and the TSS of their putative target genes. (**c-d**) AUROC and AUPRC between cis-eQTL pairs and negative control sets. See **Supplementary Methods** for how the control sets selected.

**Figure S7.**
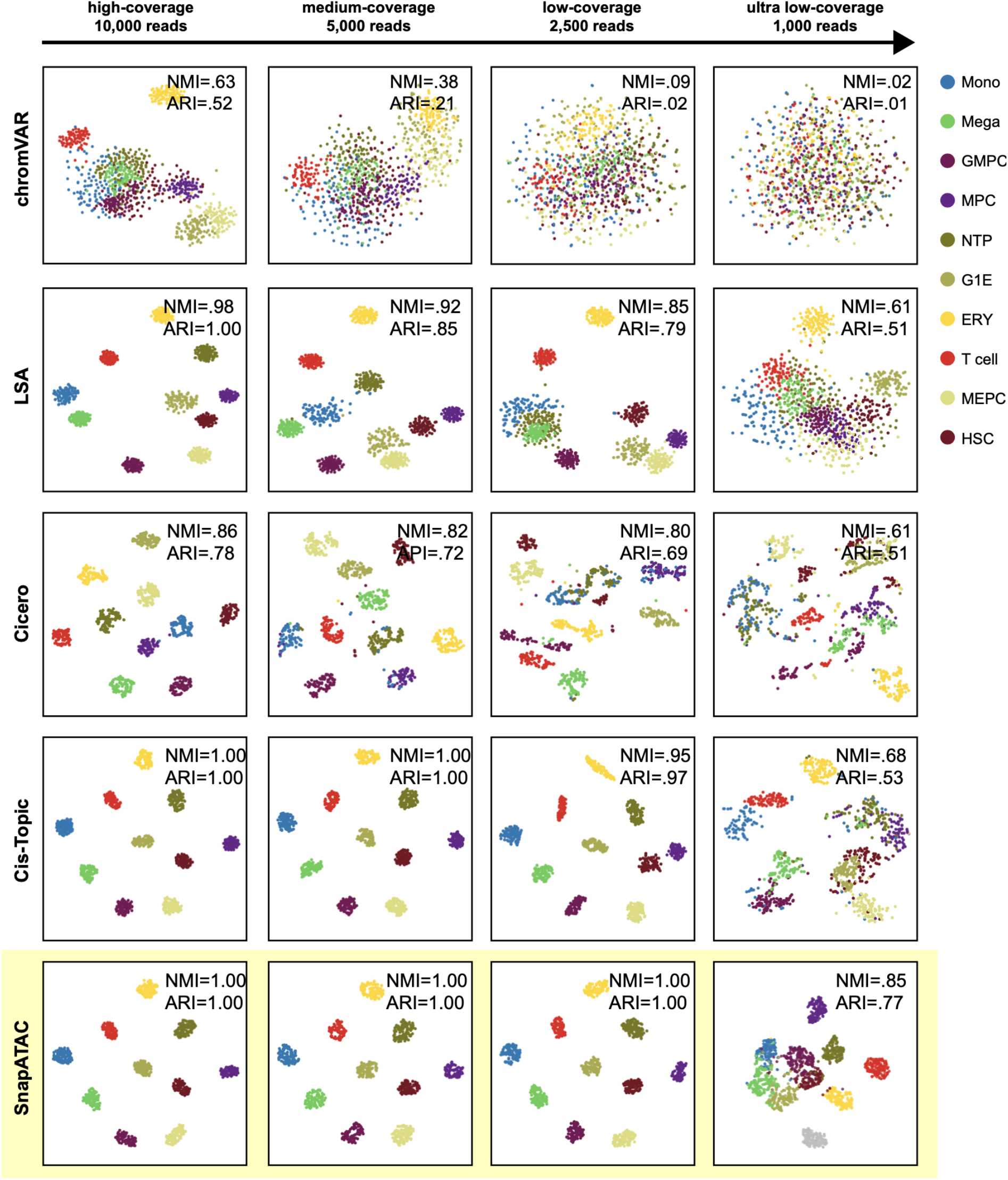
Evaluation of clustering accuracy of SnapATAC relative to alternative methods on simulated datasets. T-SNE visualization of clustering results on 1,000 simulated cells sampled from 10 bulk ATAC-seq datasets (see **Supplementary Methods** for the simulation) analyzed by five different methods – chromVAR^14^, LSA^8^, Cicero^17^, Cis-Topic^15^ and SnapATAC. Clustering results are compared to the original cell type label and the accuracy is estimated using Normalized Mutual Index (nmi). Mono: monocyte; Mega: megakaryocyte; GMPC: granulocyte monocyte progenitor cell; MPC: megakaryocyte progenitor cell; NPT: neutrophil; G1E: G1E; T cell: regulatory T cell; MEPC: megakaryocyte-erythroid progenitor cell; HSC: hematopoietic stem cell.

**Figure S8.**
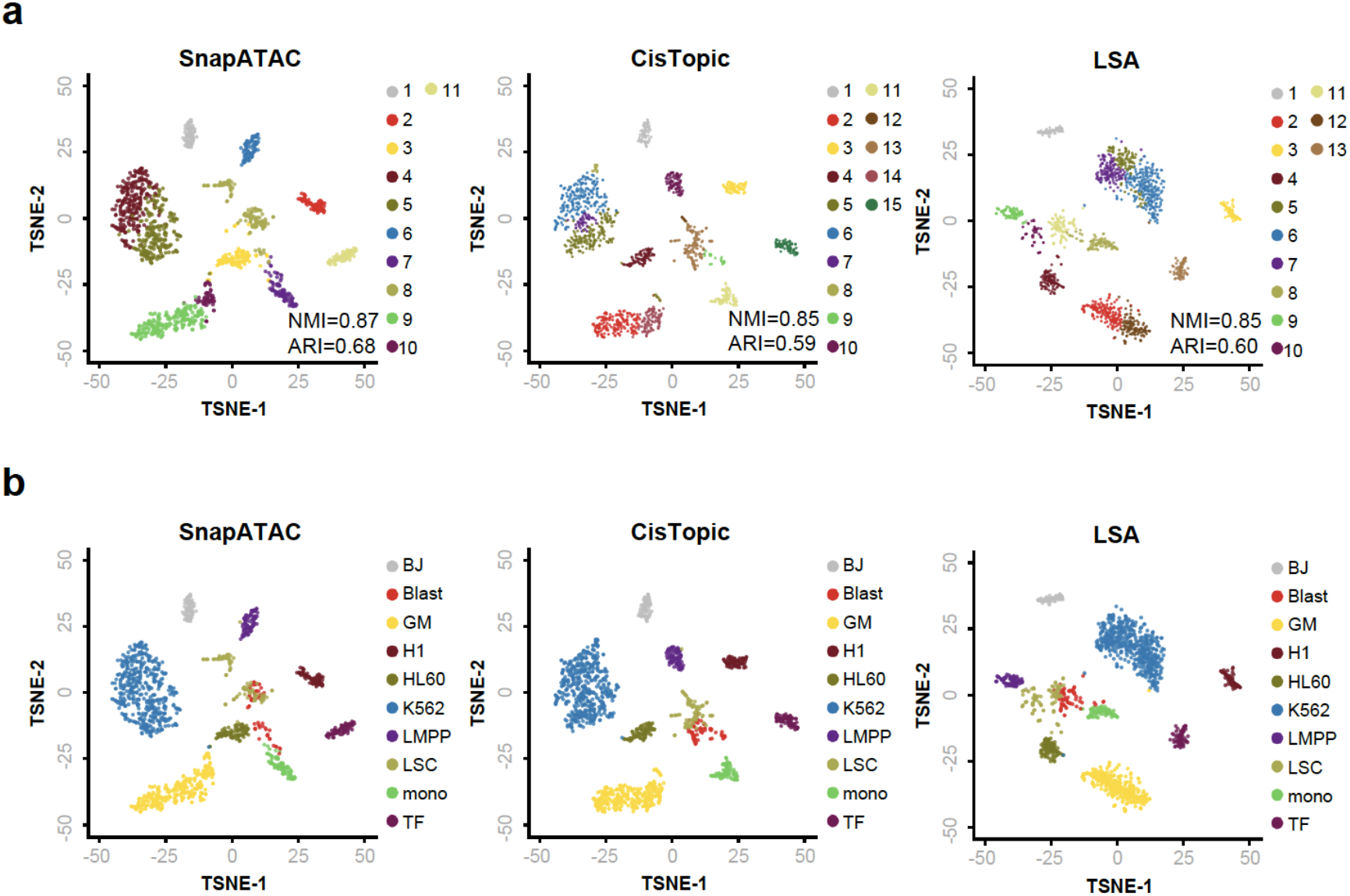
Evaluation of clustering accuracy relative to alternative methods on published single cell ATAC-seq datasets. SnapATAC (left), CisTopic (middle) and LSA (right) clustering performance on single cell ATAC-seq dataset from ten human cell lines generated using Fluidigm C1 platform^10,14^. (**a**) Clustering results are visualized using t-SNE and cells are colored by cluster labels identified by each of analysis methods. (**b**) T-SNE visualization of the human cells colored by the cell type labels. Clustering accuracy of each method is estimated by comparing the predicted clustering labels to the cell type labels. Blast: acute myeloid leukemia blast cells; LSC: acute myeloid leukemia leukemic stem cells; LMPP: lymphoid-primed multipotent progenitors; Mono: monocyte; HL60: HL-60 promyeloblast cell line; TF1: TF-1 erythroblast cell line; GM: GM12878 lymphoblastoid cell line; BJ: human fibroblast cell line; H1: H1 human embryonic stem cell line.

**Figure S9.**
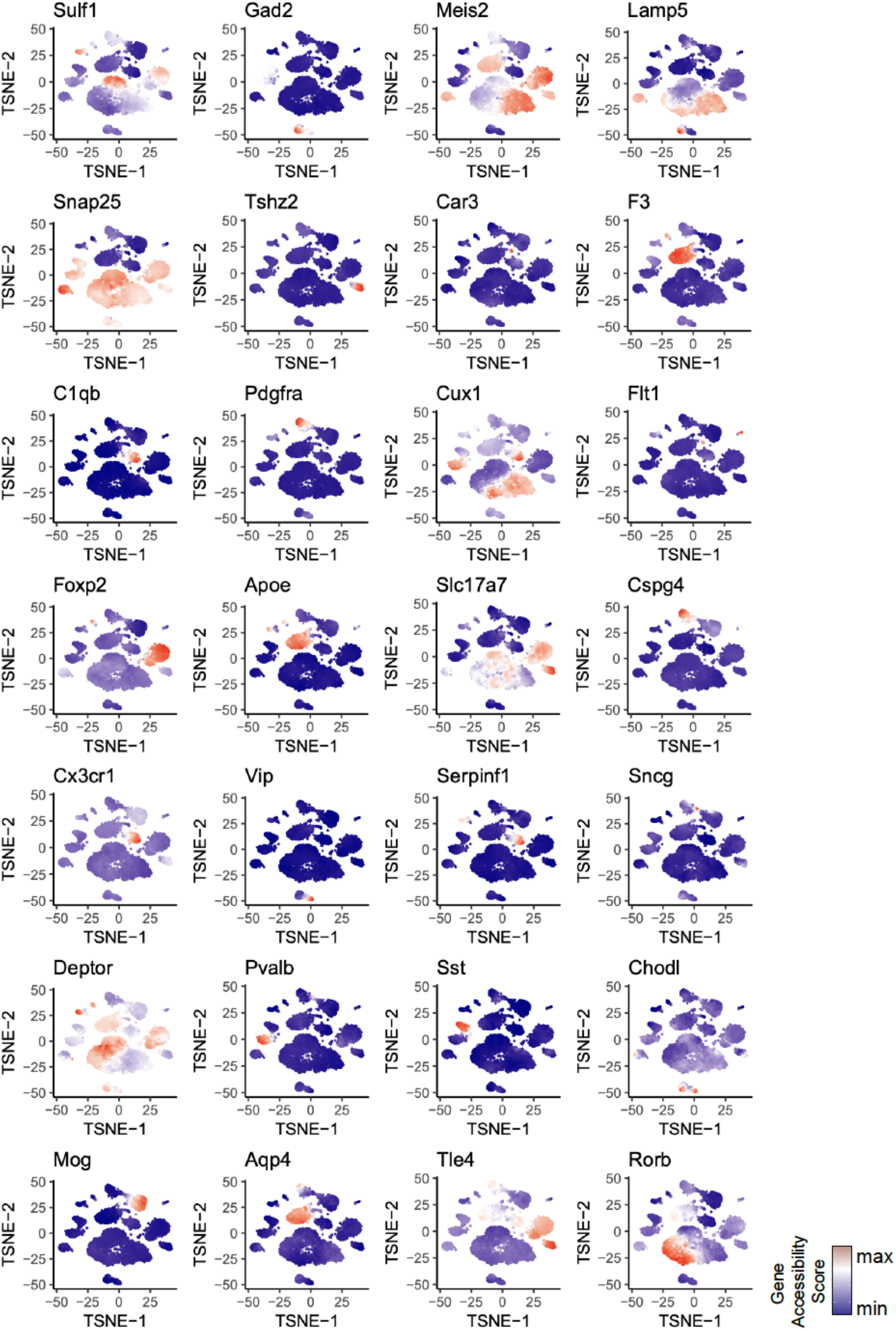
Gene accessibility score of canonical marker genes projected onto t-SNE embedding for snATAC-seq dataset from mouse secondary motor cortex. T-SNE is generated using SnapATAC; cell type specific marker genes were defined from previous single cell transcriptomic analysis in the adult mouse brain^38^; gene accessibility score is calculated using SnapATAC (**Supplementary Methods**). Data source is listed in **Supplementary Table S1**.

**Figure S10.**
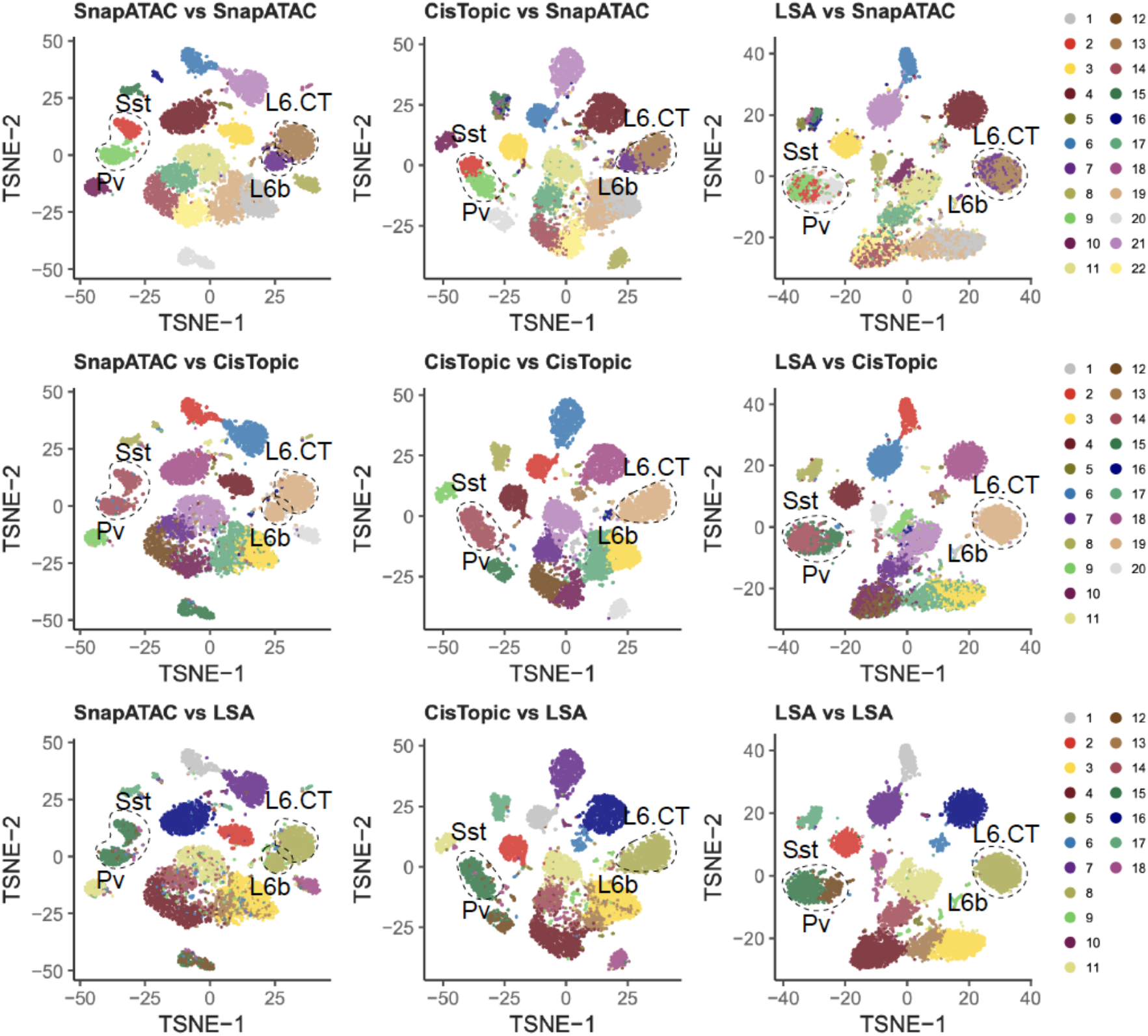
Evaluation of clustering sensitivity of SnapATAC relative to alternative methods on mouse secondary motor cortex snATAC-seq. Three methods (cisTopic, LSA and SnapATAC) were used to analyze a dataset that contained ∼10k single nucleus ATAC-seq profiles from the mouse secondary motor cortex. Pairwise comparison of the clustering results is shown by projecting the cluster label identified using one method onto the t-SNE visualization generated by another method (cluster vs. visualization). Black dash line circles highlight the rare pollutions (Sst, Pv, L6b and L6.CT) that were only identified by SnapATAC. Data source is listed in **Supplementary Table S1**.

**Figure S11.**
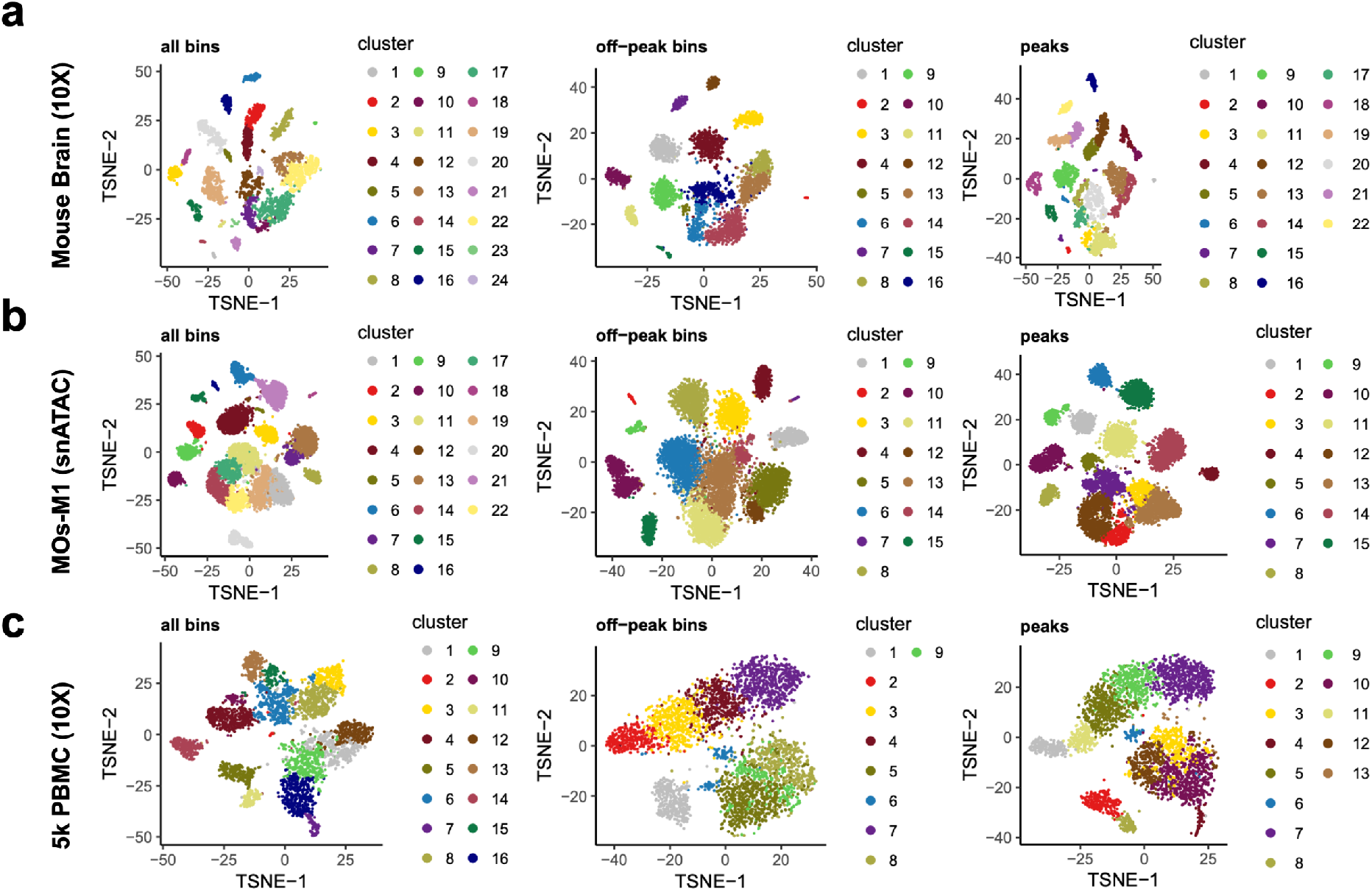
Off-peak reads distinguish major cell types in heterogenous samples. (**a-c**) SnapATAC clustering result on three benchmarking datasets using all bins versus clustering result only using bins that are not overlapped with peaks. Data source is listed in **Supplementary Table S1**.

**Figure S12.**
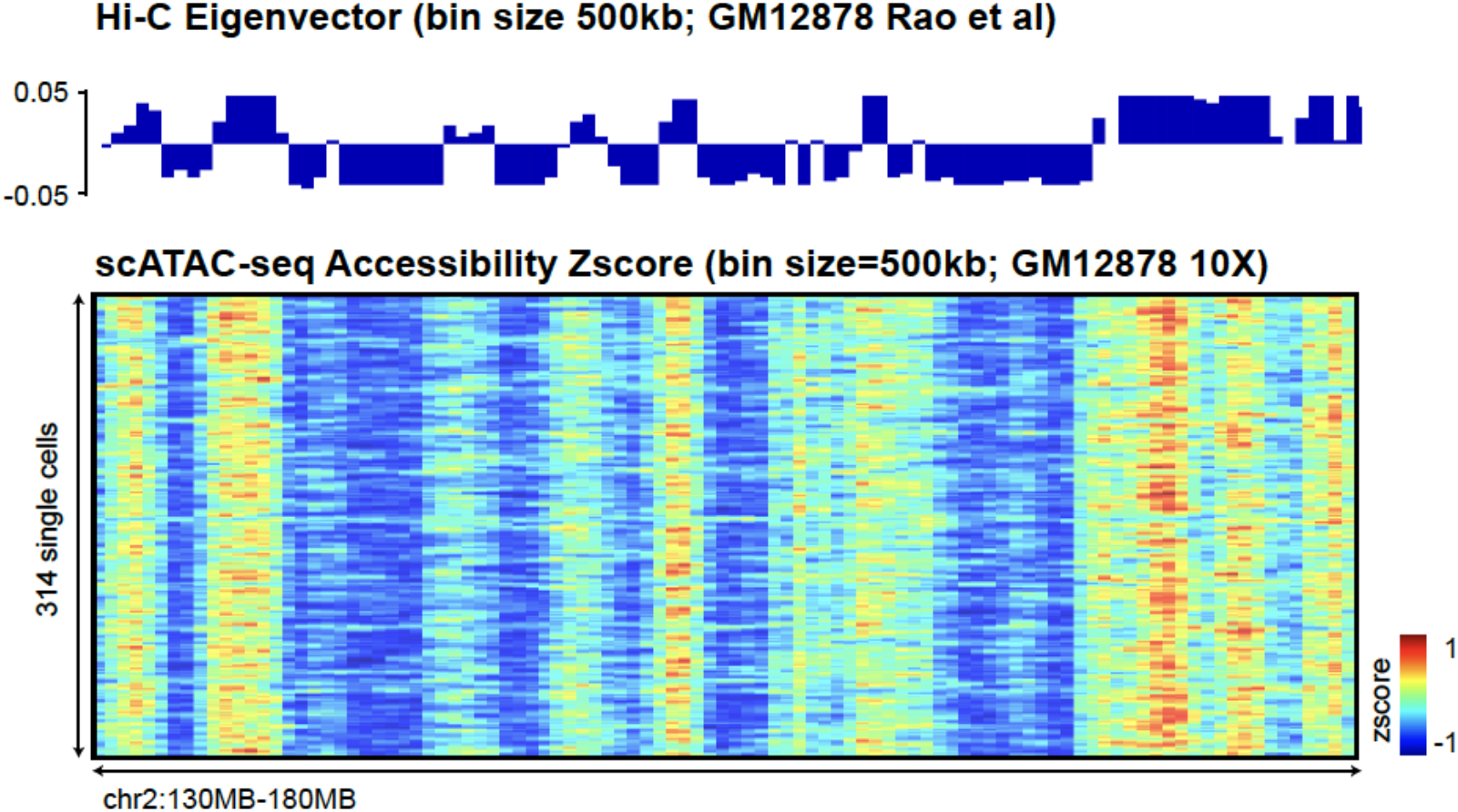
Off-peak reads reflect higher-order chromatin structure. At 500kb bin resolution, profile of compartments identified using Hi-C^58^ in GM12878 overlaid the density of “off-peak” reads for 314 cells from GM12878 10X scATAC-seq library. Data source is listed in **Supplementary Table S1**.

**Figure S13.**
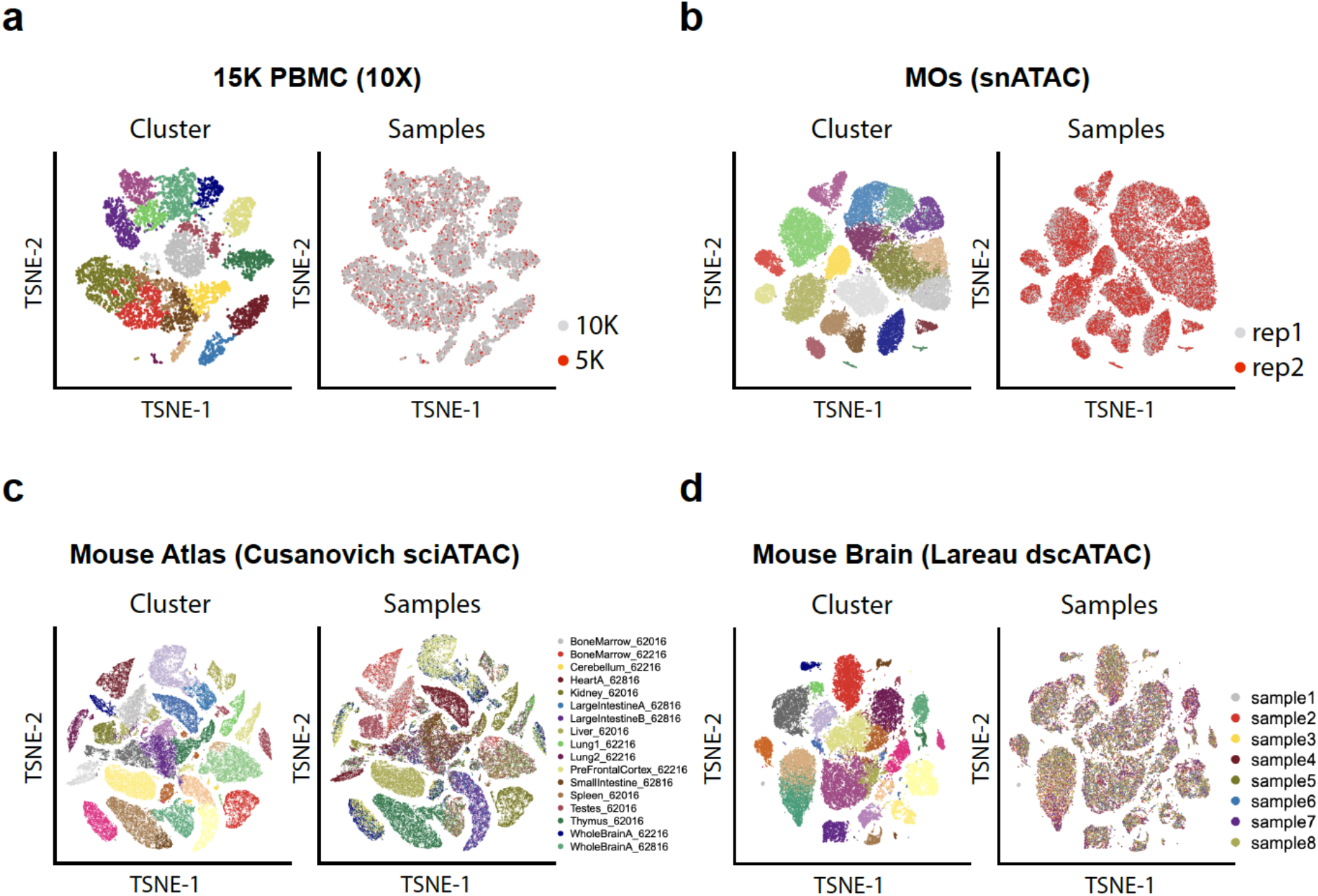
SnapATAC is robust to technical variation. Two-dimensional t-SNE visualization of four benchmarking datasets generated using SnapATAC. Cells are color by cluster label (left) and sample label (right). (**a**) 15k PBMC (10X) – a combination of two datasets (PBMC 5k and 10k) publicly available from 10X genomics. (**b**) MOs (snATAC) – an in-house dataset that contains two biological replicates from secondary motor cortex in the adult mouse brain generated using single nucleus ATAC-seq. (**c**) Mouse Atlas (Cusanovich 2018) – a published dataset that contains over 80K cells from 13 different mouse tissues generated using multiplexing single cell ATAC-seq. (**d**) Mouse Brain (Lareau dscATAC) – a published dataset that contains 46,652 cells from 8 samples in the adult mouse brain generated using BioRad droplet-based single cell ATAC-seq. Data source is listed in **Supplementary Table S1**.

**Figure S14.**
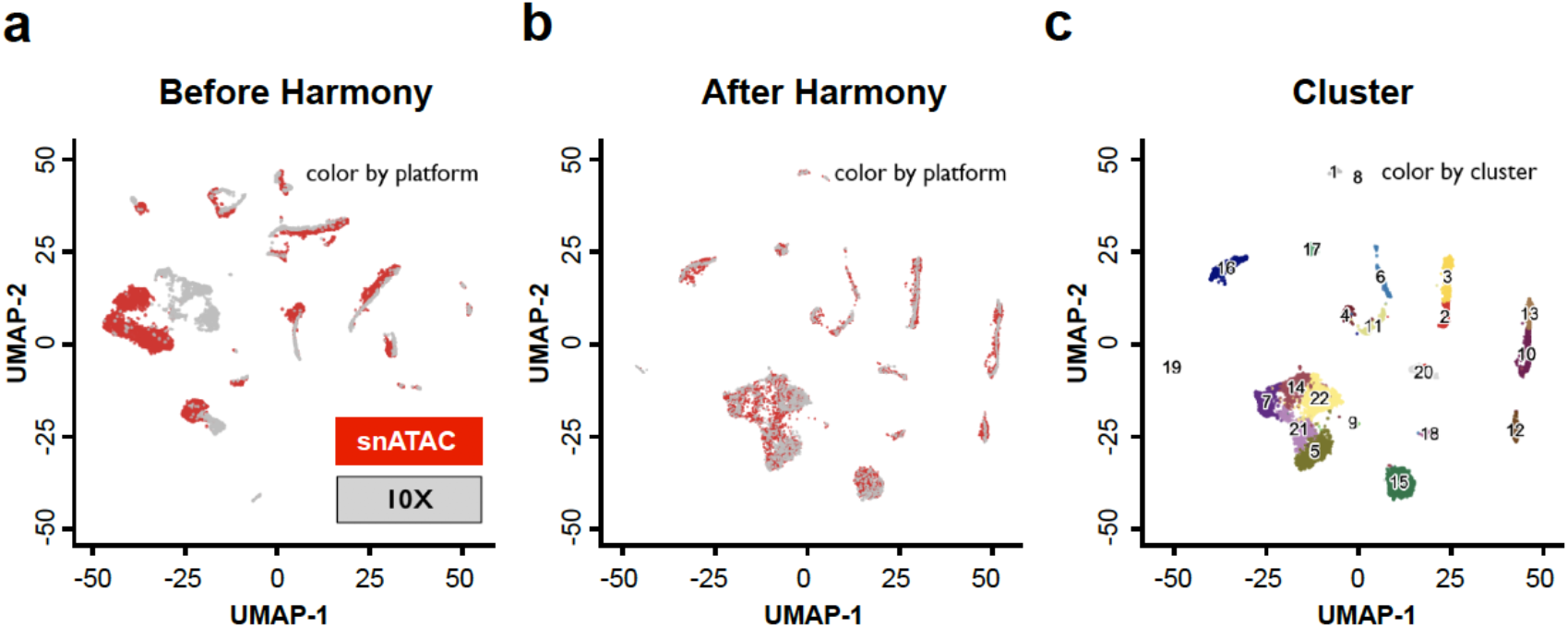
SnapATAC eliminates batch effect using Harmony^23^. The joint UMAP visualization of two datasets of mouse brain generated using combinatorial indexing single nucleus ATAC-seq (MOs-M1 snATAC) and droplet-based platform (Mouse Brain 10X) before (**a**) and after (**b**) performing batch effect correction using Harmony. Data source is listed in **Supplementary Table S1**.

**Figure S15.**
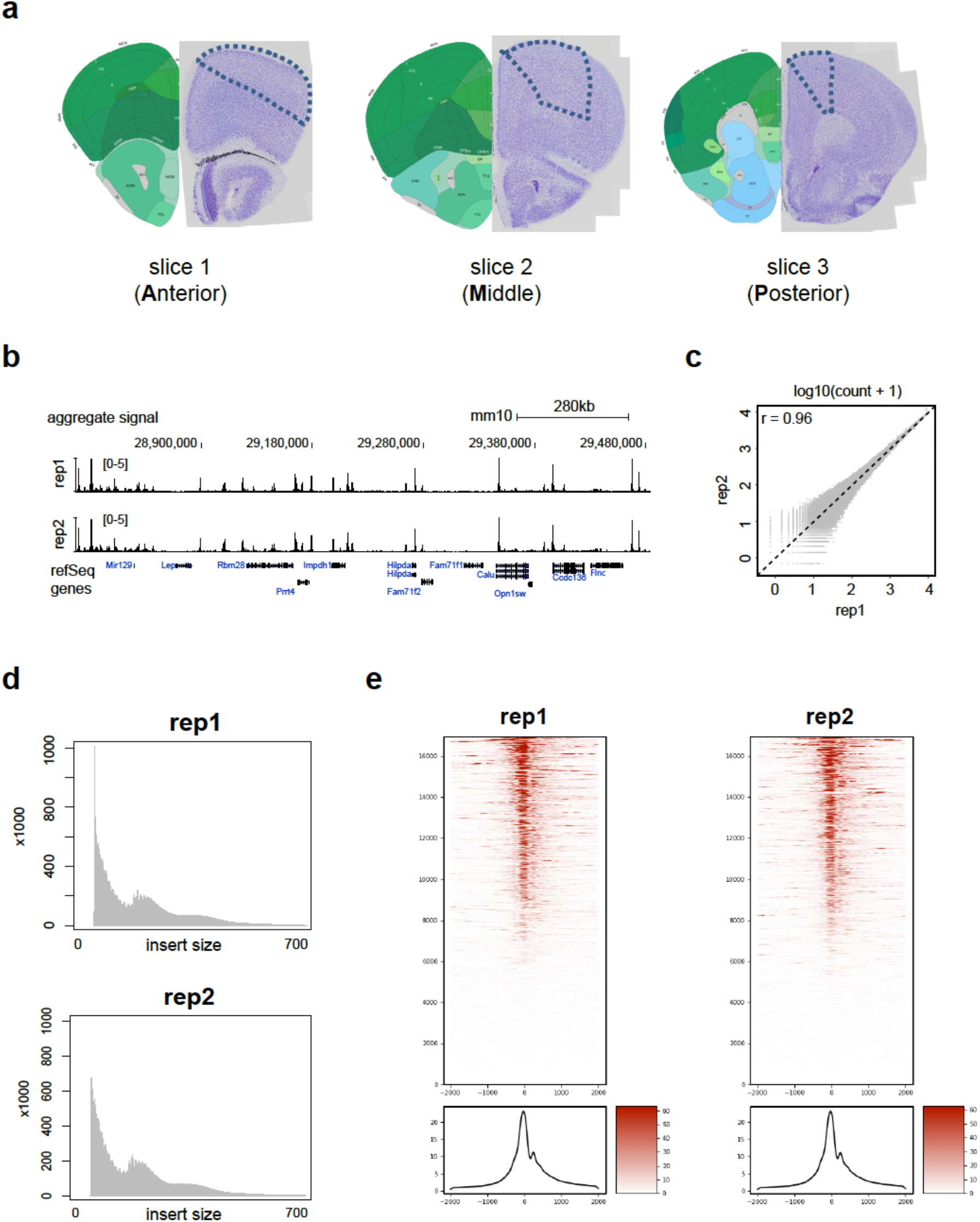
Single nucleus ATAC-seq datasets are reproducible between biological replicates. (**a**) Illustration of dissection. Posterior view of three 0.6 mm coronal slices from which the secondary motor cortex (MOs) was dissected. The right side on each image depicts the corresponding view from the Allen Brain Atlas. The left side correspond to the Nissl staining of the posterior side of each slice. The MOs region was manually dissected according to the dashed lines on each slice and following the MOs as depicted in plates 27, 33, and 39 of the Allen Brain Atlas (left side images in figure). Each slice contains two biological replicates named as A1, A2, M1, M2, P1 and P2 (A: Anterior; M: Middle; P: Posterior). In this study, A1, M1 and P1 is combined as replicate 1 and A2, M2 and P2 are combined as replicate 2. (**b**) Genome-browser view of aggregate signal for two biological replicates. (**c**) Pearson correlation of count per million (CPM) at peaks between two replicates. (**d**) Insert size distribution and (**e**) TSS enrichment score for two biological replicates.

**Figure S16.**
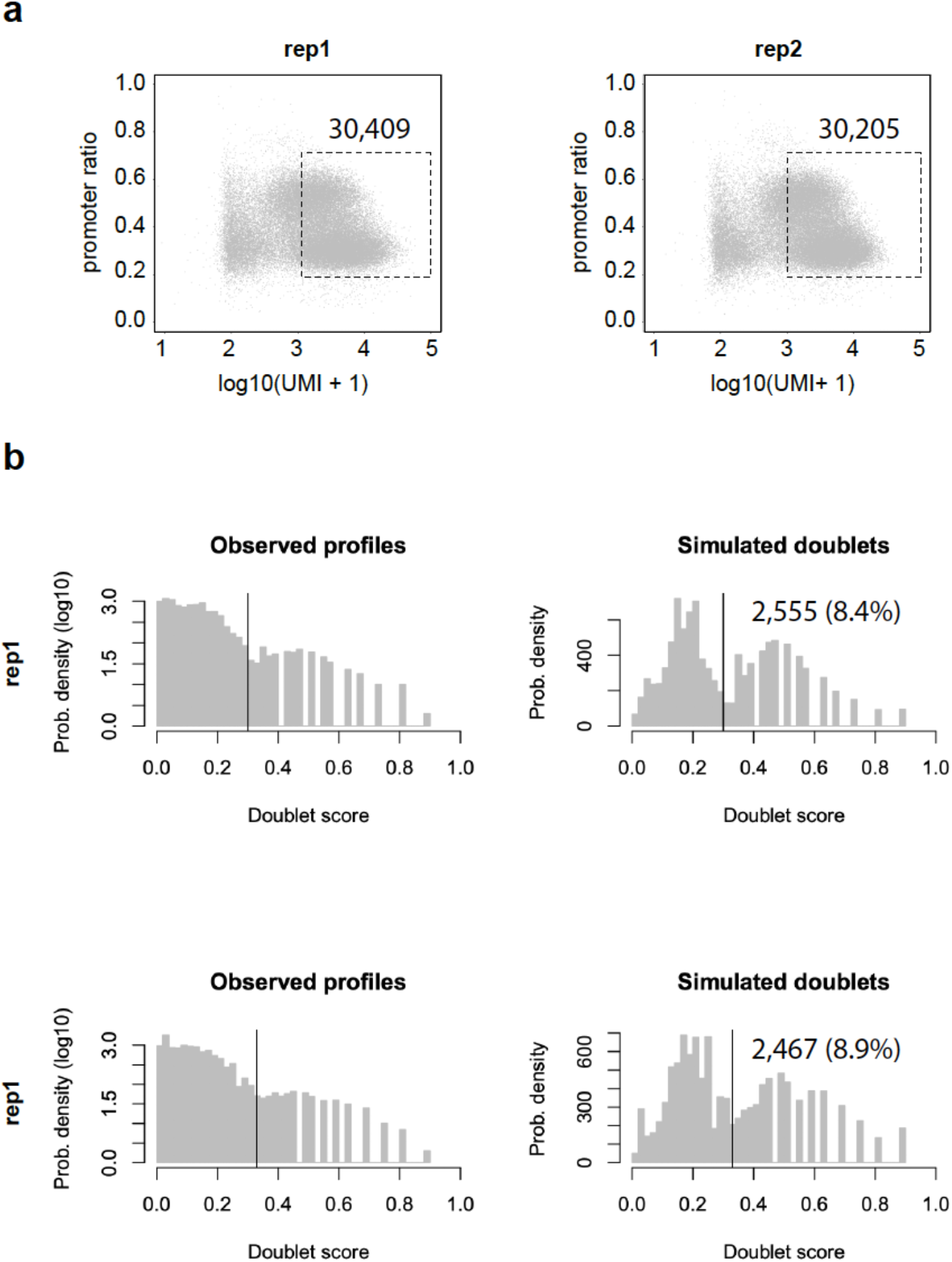
Barcode selection of MOs. (**a**) Cells of unique fragments within the range of 1,000-100,000 and fragments in promoter ratio within the range of 0.2-0.7 were selected. This resulted in 30,409 and 30,205 nuclei for two replicates. (**b**) With 5kb cell-by-bin matrix as input matrix, putative doublets were identified using Scrublets^37^, which predicted 2,555 (8.4%) and 2,467 (8.9%) nuclei to be doublets for each replicate. The predicted doublet ratio is similar to the theoretical calculation of doublet ratio for multiplexing single cell ATAC-seq experiment^5,7^.

**Figure S17.**
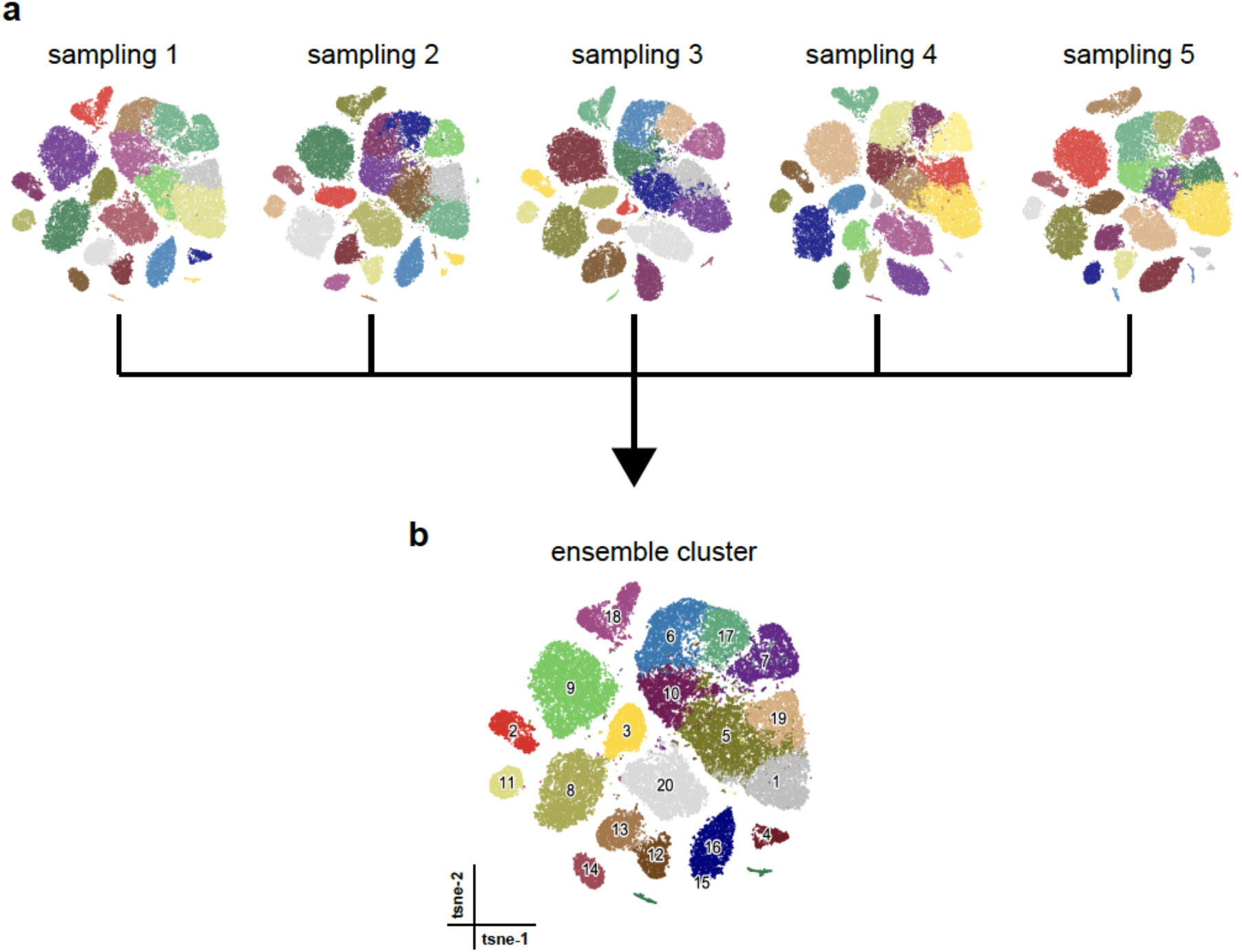
Consensus clustering of MOs. (**a**) Five clustering results were generated using SnapATAC with different set of landmarks (10,000). (**b**) These five clustering solutions were combined to create a consensus clustering which identified 20 clusters in MOs (**Supplementary Methods**).

**Figure S18.**
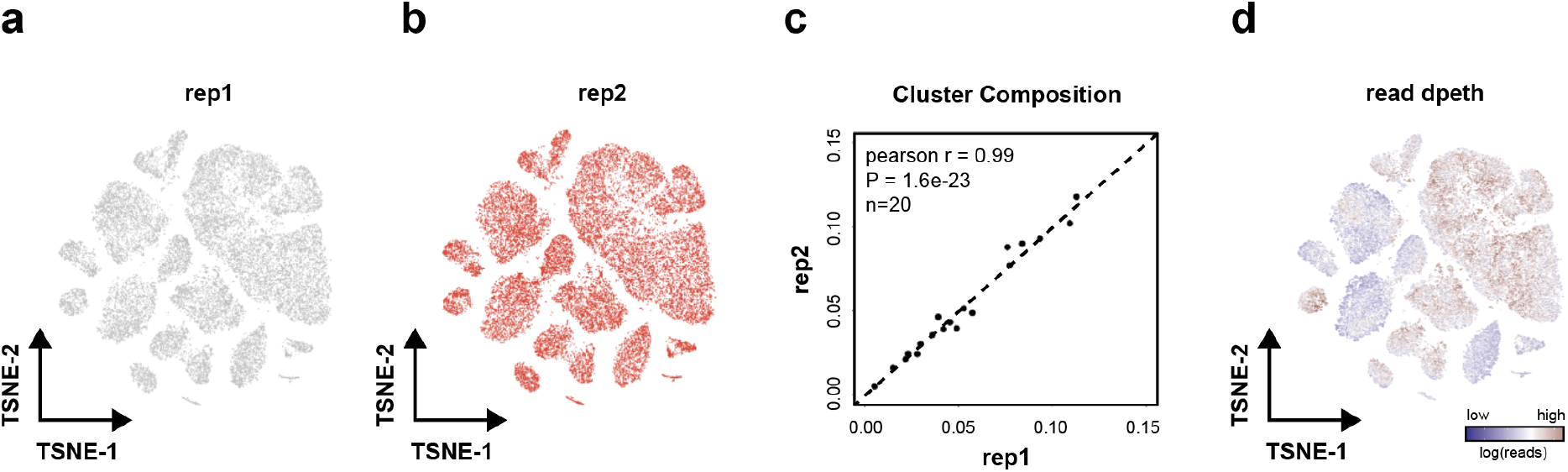
MOs clustering result is reproducible between biological replicates. (**a-b**) T-SNE visualization of cells from two biological replicates. (**c**) The cluster composition is highly reproducible between two biological replicates (r=0.99; P-value = 1.6e-23); (**d**) T-SNE visualization of cells with color scaled by sequencing depth.

**Figure S19.**
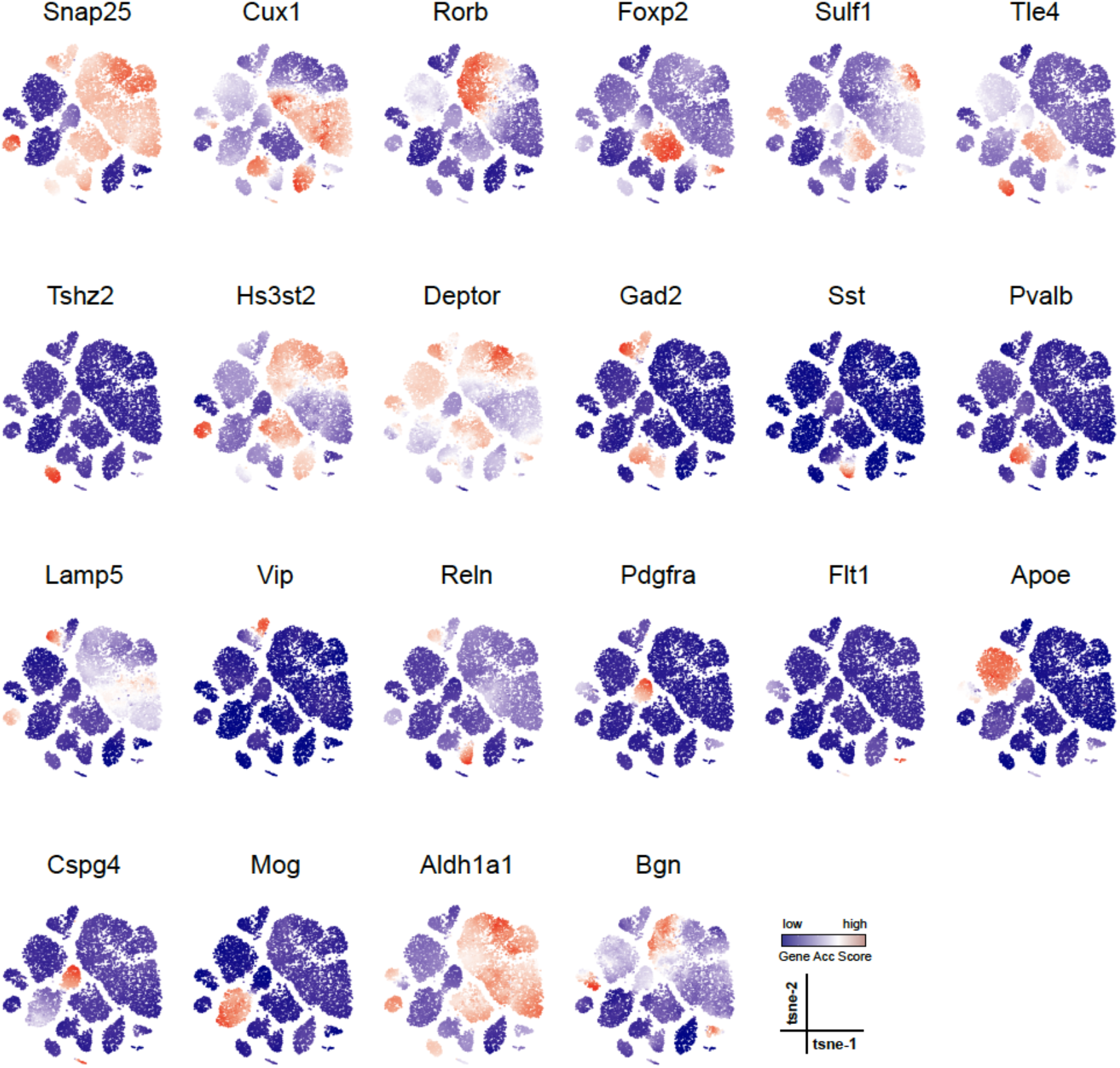
Gene accessibility score of canonical marker genes projected onto MOs t-SNE embedding to guide the cluster annotation. T-SNE is generated using SnapATAC for MOs; cell type specific marker genes was defined from previous single cell transcriptomic analysis in adult mouse brain^38^; gene accessibility score is calculated using SnapATAC (**Supplementary Methods**) and projected to the t-SNE embedding.

**Figure S20.**
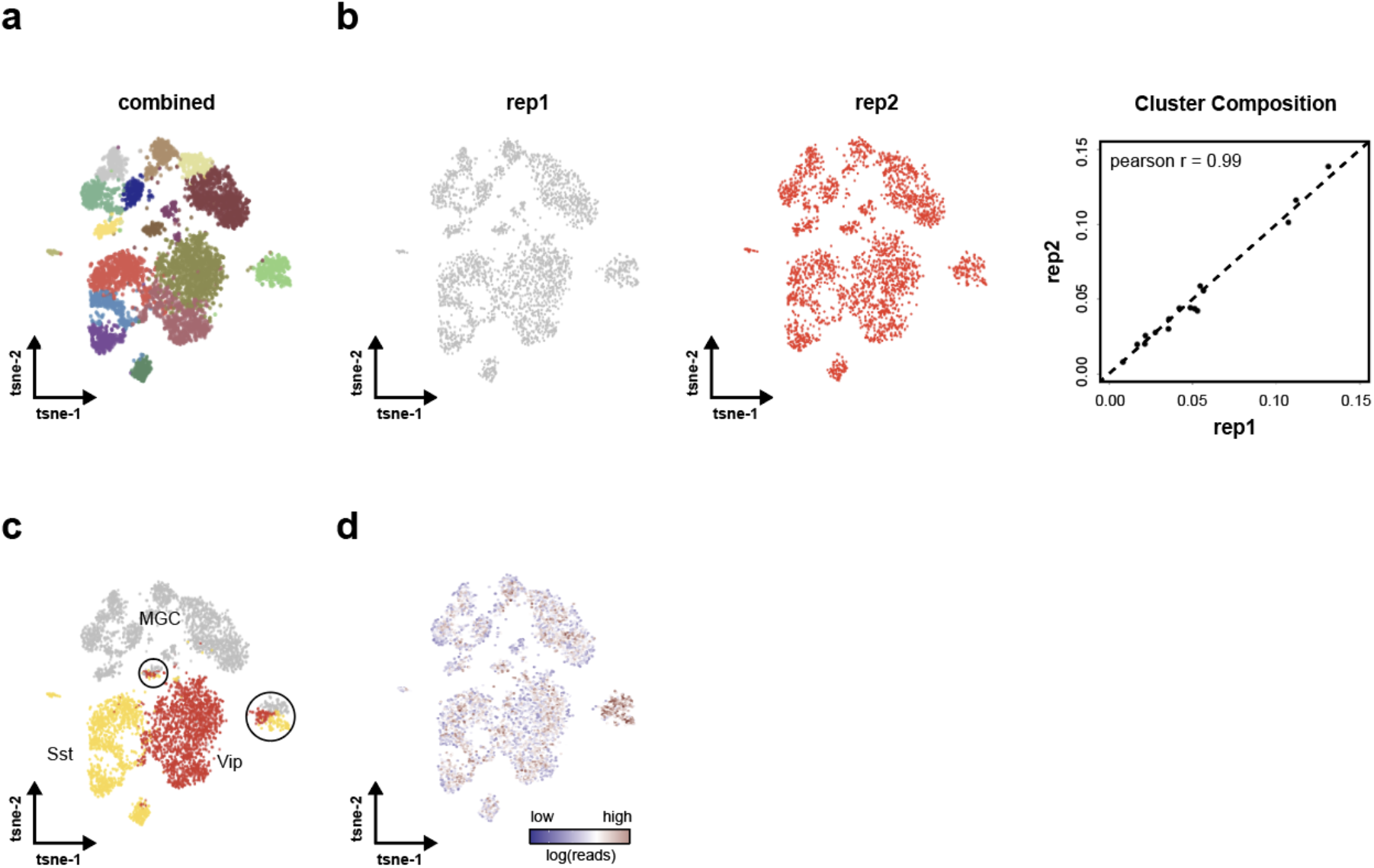
Iterative clustering identifies 17 GABAergic neuronal subtypes. (**a**) Sub-clustering of 5,940 GABAergic neurons identified 17 distinct cell clusters. (**b**) Cluster composition was highly reproducible between two biological replicates. (**c**) TSNE visualization of 5,940 GABAergic neurons colored by cell types identified in the initial clustering (shown in **Fig. 5a**). Black circles mark clusters that are potential doublets, a mixture of multiple cell types. (**d**) TSNE plot of GABAergic neurons colored by sequencing depth.

**Figure S21.**
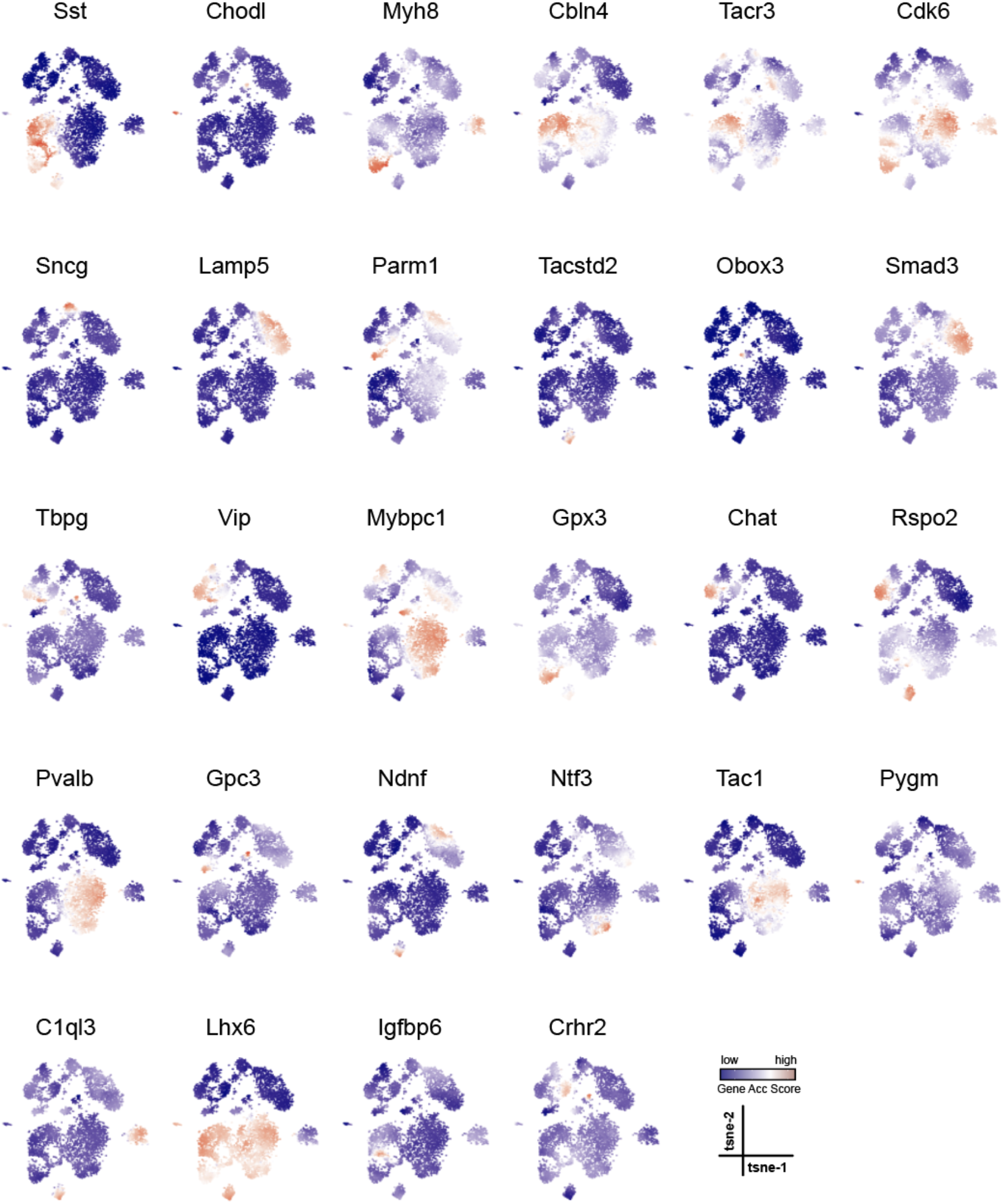
Gene accessibility score of marker genes projected onto t-SNE embedding from GABAergic neurons to guide the cluster annotation. Iterative clustering is performed against GABAergic neurons to identify subtypes. Twenty eight cell type specific marker genes were defined from previous single cell transcriptomic analysis in adult mouse brain^38^; gene accessibility score is calculated using SnapATAC (**Supplementary Methods**).

**Figure S22.**
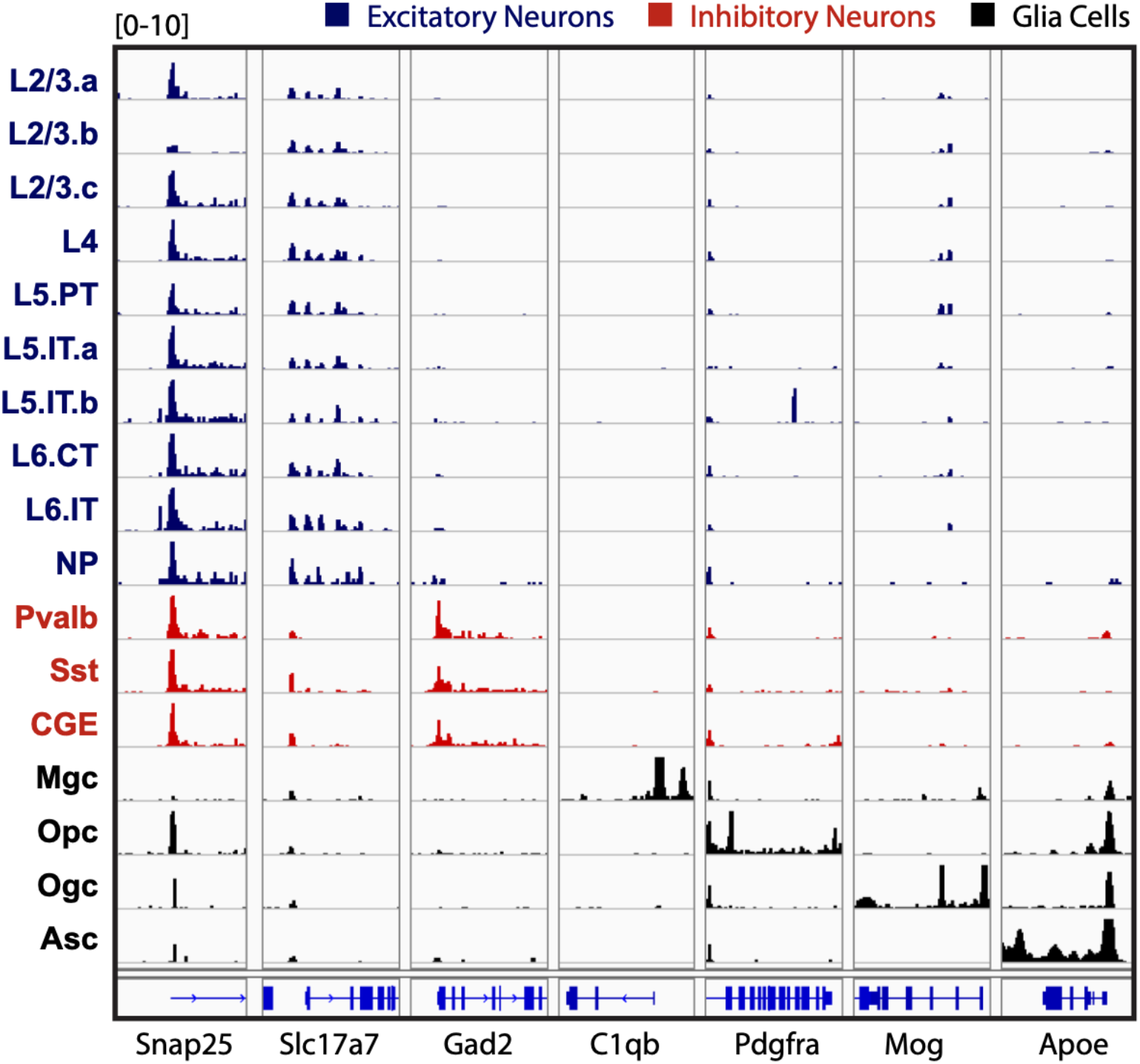
Genome browser view of aggregate signal for each of the major cell populations identified in the adult mouse brain (Fig. 5a).

**Figure S23.**
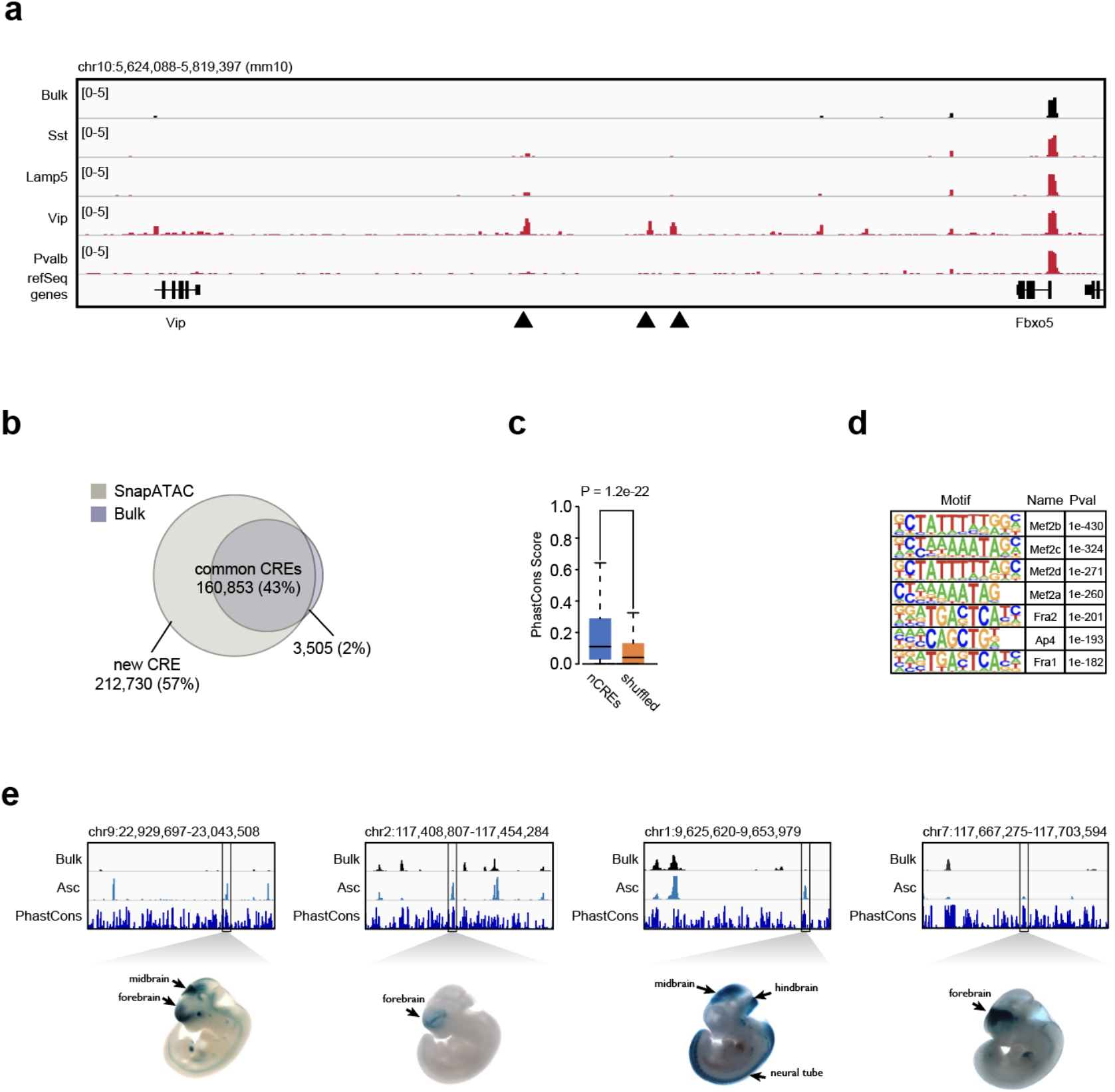
SnapATAC uncovers novel candidate *cis*-regulatory elements in rare cell types. (**a**) Genome browser view of 20Mb region flanking gene *Vip*. Dash line highlight five regulatory elements specific to Vip subtypes that are under-represented in the conventional bulk ATAC-seq signal. (**b**) Over fifty percent of the regulatory elements identified from 20 major cell populations are not detected from bulk ATAC-seq data. (**c**) Sequence conservation comparison between the new elements and randomly chosen genomic regions. (**d**) Top seven motifs enriched in Pv-specific new elements. (**e**) Examples of four new elements that were previously tested positive in transgenic mouse assays (from VISTA database). Bulk: Bulk ATAC-seq; Asc: aggregated signal from astrocyte population (ASC) in the adult mouse brain as shown in **Fig. 5a**.

**Figure S24.**
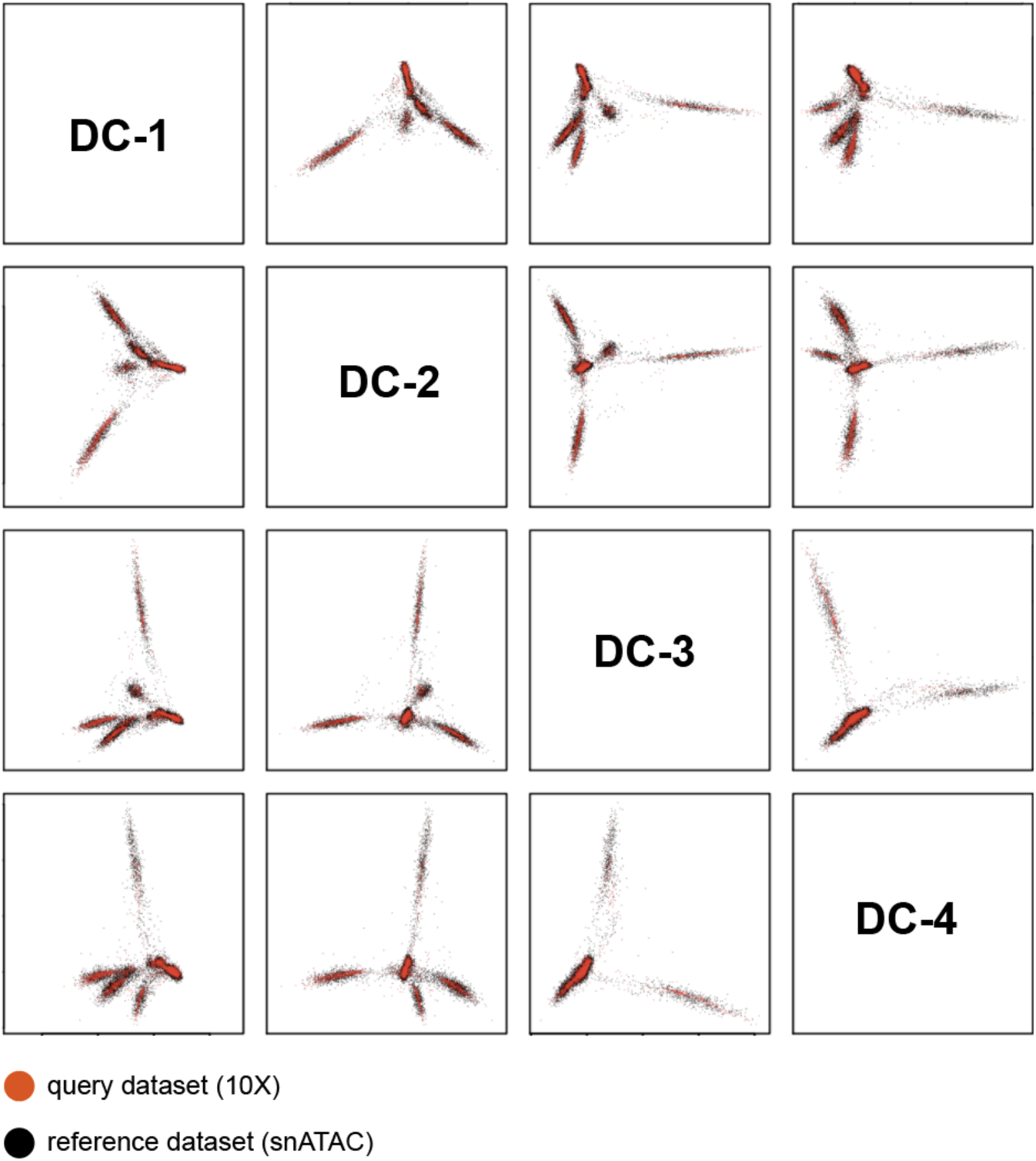
Joint embedding for query (Mouse Brain 10X) and reference dataset (MOs snATAC). The query dataset (10X) is projected onto the low dimension embedding space precomputed for the reference dataset (snATAC). Batch effect is corrected using Harmony. Pairwise plot of the first four dimentions in which cells are colored by dataset - red for query cells (Mouse Brain 10X) and black for reference cells (MOs snATAC). Data source as listed in **Supplementary Table S1**.

**Figure S25.**
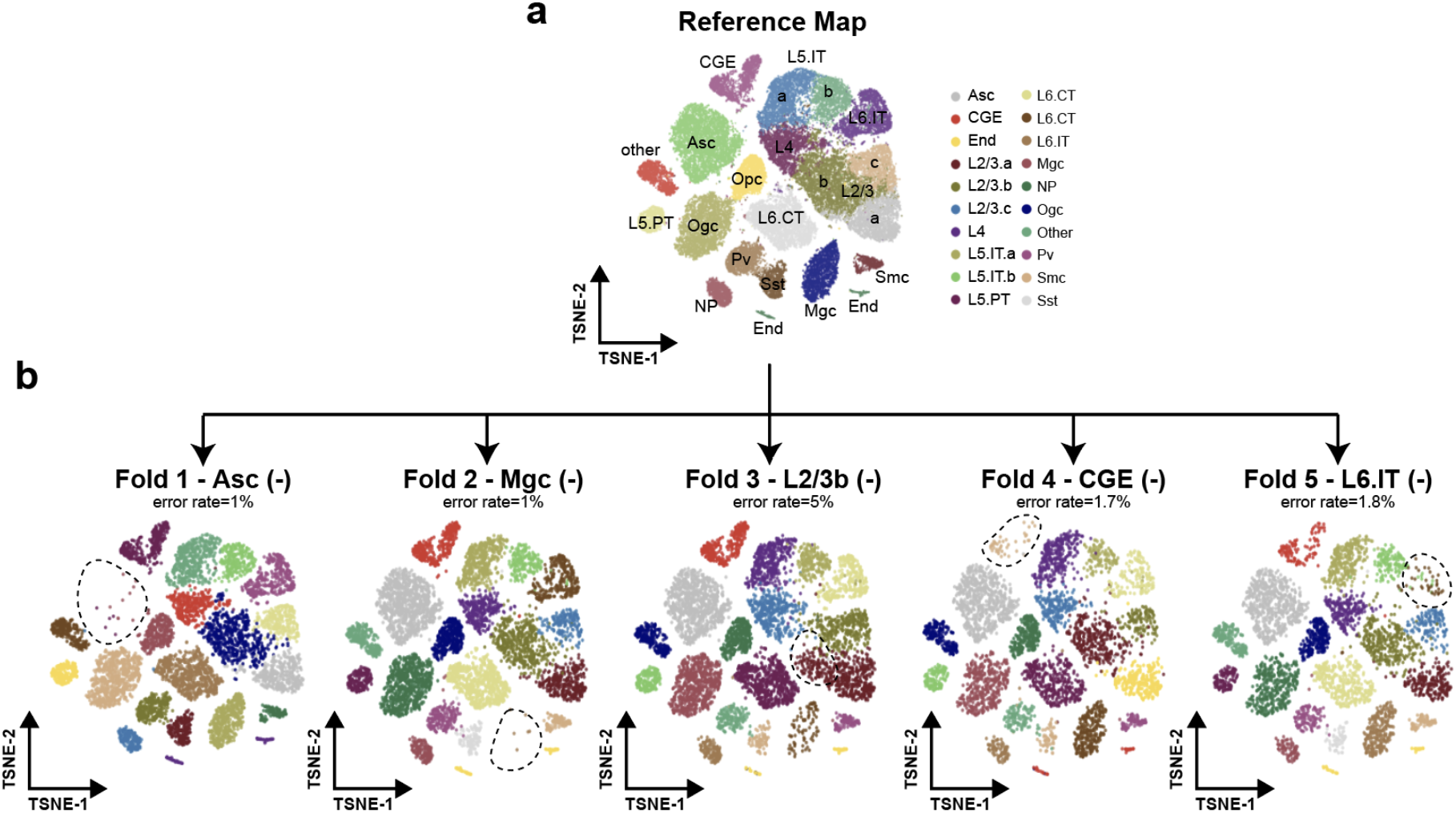
SnapATAC is robust for supervised annotation of datasets containing cell types missing in the reference atlas. (**a**) Two-dimensional t-SNE visualization of the reference dataset MOs (snATAC). (**b**) A five-fold cross validation is performed to this reference dataset. For each fold, we introduce perturbation to the 80% training dataset by randomly dropping one cell type (Asc, Mgc, L2/3b, CGE and L6.IT). We then predict on the 20% test dataset using the model learned from the perturbed training dataset. The prediction accuracy for each fold is shown in (**b**) and cell type removed from the training dataset are highlighted by the dash-line circles.

**Figure S26.**
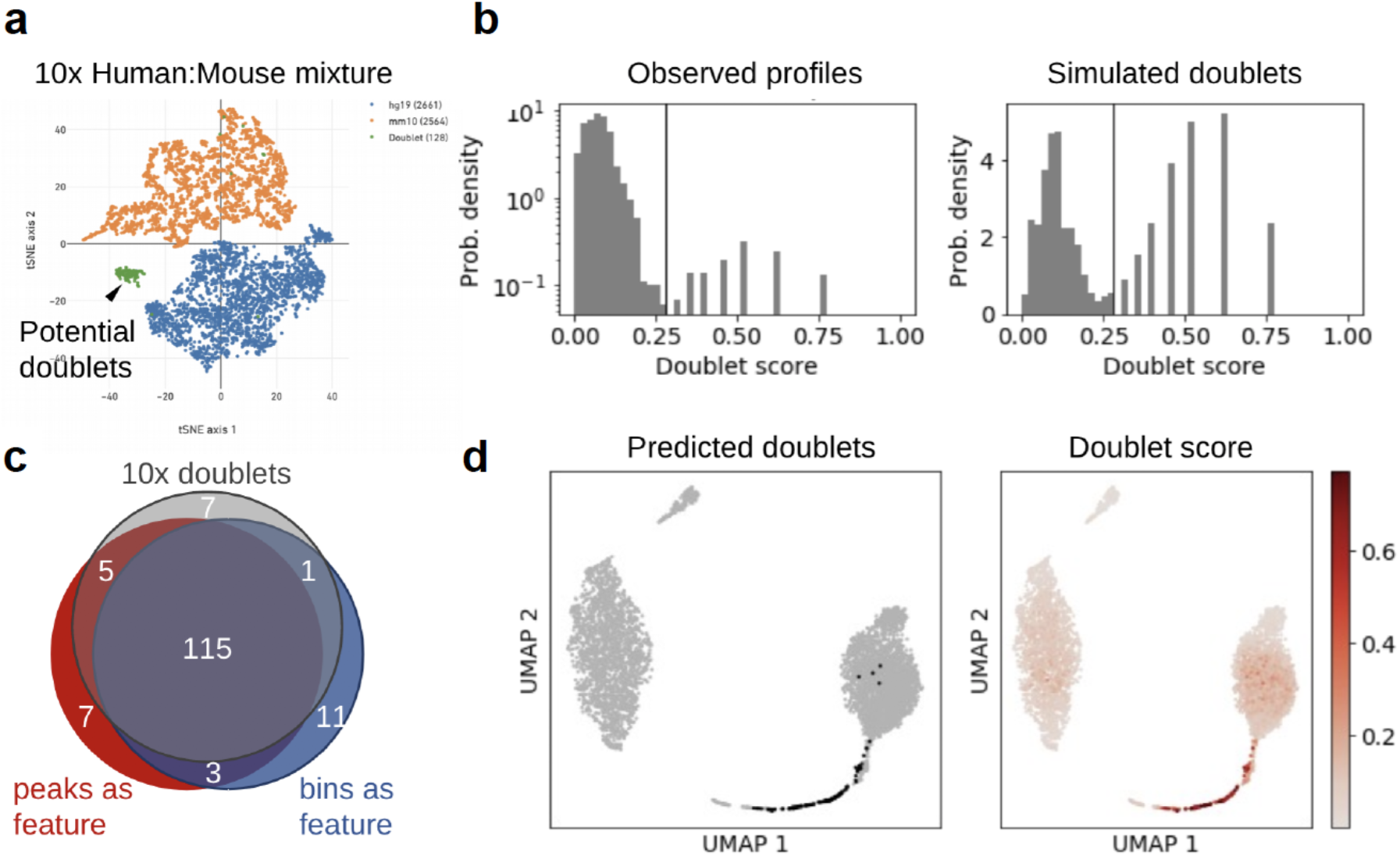
Doublets detection using Scrublet. (**a**) T-SNE representation of a dataset (hgmm_1k 10X) that contained 1,000 human (GM12878) and mouse (A20) cells. Cells are colored by species determined based on the alignment ratio between human and mouse genome. Orange: A20; blue: GM12878; green: putative doublets. (**b**) Distribution of doublet score for putative doublets and simulated doublets estimated using Scrublet^37^. (**c**) Doublets are predicted using cell-by-peak and cell-by-bin matrix separately. Venn diagram show the overlap between Scrublet-predicted doublets using peak or bin matrix and doublets identified based on alignment ratio. (**d**) Doublets scores projected onto the UMAP embedding.

